# Bilirubin is a new ligand for nuclear receptor Liver Receptor Homolog-1

**DOI:** 10.1101/2024.05.05.592606

**Authors:** Pratima Chapagain, Zeinab Haratipour, M. Merced Malabanan, Woong Jae Choi, Raymond D. Blind

**Author notes:** Equal Contributions.

## Abstract

The nuclear receptor Liver Receptor Homolog-1 (LRH-1, *NR5A2*) binds to phospholipids that regulate important LRH-1 functions in the liver. A recent compound screen unexpectedly identified bilirubin, the product of liver heme metabolism, as a possible ligand for LRH-1. Here, we show unconjugated bilirubin directly binds LRH-1 with apparent *K_d_*=9.3uM, altering LRH-1 interaction with all transcriptional coregulator peptides tested. Bilirubin decreased LRH-1 protease sensitivity, consistent with MD simulations predicting bilirubin stably binds LRH-1 within the canonical ligand binding site. Bilirubin activated a luciferase reporter specific for LRH-1, dependent on co-expression with the bilirubin membrane transporter *SLCO1B1*, but bilirubin failed to activate ligand-binding genetic mutants of LRH-1. Gene profiling in HepG2 cells shows bilirubin selectively regulated transcripts from endogenous LRH-1 ChIP-seq target genes, which was significantly attenuated by either genetic knockdown of LRH-1, or by a specific chemical competitor of LRH-1. Gene set enrichment suggests bilirubin and LRH-1 share roles in cholesterol metabolism and lipid efflux, thus we propose a new role for LRH-1 in directly sensing intracellular levels of bilirubin.

## Introduction

Nuclear receptors are a family of transcription factors regulated by hydrophobic lipid ligands such as cholesterol-based bile acids, steroids^1^, fatty acids^2,3^, phospholipids^4^ and heme^5^, with important functions conserved throughout all metazoans^6,7^. Liver Receptor Homolog-1 (LRH-1, *NR5A2*) is a key nuclear receptor in the development^8–12^ and adult function^13–17^ of the liver^18–22^, pancreas^18,23,24^ and gut^9,25–29^, with critical roles in bile acid and lipid metabolism. Although LRH-1 lacks a definitively established ligand in humans or any other organism, many lines of evidence published by our group and others have suggested phospholipids regulate LRH-1 structure and function^15,30–32^ with complete physiological and structural details established for phosphatidylcholines^17,33,34^ and more limited binding and co-crystal structures for phosphatidylinositols^32,35^ and phosphatidylethanolamines^15,32,36^. Phospholipids bind the canonical site in the ligand-binding domain of LRH-1 to allosterically regulate LRH-1 conformation, which serves to alter LRH-1 interaction with transcriptional coregulators and LRH-1 functions in transcription, akin to other nuclear receptors such as the close homolog Steroidogenic Factor-1 (SF-1, *NR5A1*)^31,33,34,37–42^. The unusually wide array of phospholipid ligands of LRH-1 are accommodated by the relatively large ligand-binding pocket of human LRH-1, so it remains unclear if phospholipids are the only natural ligands capable of directly regulating LRH-1.

We fortuitously identified unconjugated bilirubin as a potential ligand for LRH-1 in a recent chemical compound screen^43^. Bilirubin is a breakdown product of heme present in the liver and the precursor of glucuronidated bilirubin, an important component of bile. LRH-1 is well known to regulate bile acid metabolism^19^, and heme is known to regulate the nuclear receptor Rev-Erb^5^, however no evidence has suggested any metabolite of heme directly binds or regulates LRH-1, despite these molecules sharing roles in hepatic cholesterol metabolism^19,44^ and lipid transport^45–47^. LRH-1 is known to activate expression of hepatic bilirubin membrane transporters when ratios of unconjugated to conjugated bilirubin are high^46–48^ but the molecular details of how LRH-1 might sense these bilirubin ratios have remained almost completely uninvestigated.

Here, we present multiple, orthogonal lines of evidence suggesting unconjugated bilirubin directly regulates human LRH-1, *in vitro*, *in silico* and in cells, and that in parallel with other nuclear receptors^49–52^, human LRH-1 acts as a bilirubin sensor. Although the physiological consequences of bilirubin regulation of LRH-1 are not established here, our data introduce bilirubin as the first non-phospholipid regulatory metabolite of the nuclear receptor LRH-1, and further support the notion that bilirubin acts as a second messenger signaling molecule, capable of directly regulating transcription through receptor-mediated events within the nuclear compartment.

## Results

### Bilirubin regulates purified LRH-1 ligand binding domain functions *in vitro*

We previously identified unconjugated bilirubin (**Fig 1A**) to bind purified LRH-1 in an *in vitro* high-throughput compound screen^43^. Here, we used a FRET-based competition assay^43^ (**Fig S1A-C**) to approximate the apparent dissociation constant of unconjugated bilirubin for LRH-1 ligand binding domain (9.3µM, 95%CI = 4.6-22.0µM, **Fig 1B**), and confirmed that approximate binding in 2 other independent assays, a tryptophan fluorescence assay and a melting temperature assay (**Fig 1C, S2A-D**). A pocket mutant of LRH-1 (A349W) previously shown to attenuate ligand binding^53^, also had attenuated bilirubin binding (≥42.7µM, 95%CI = 22.7-89.6µM) (**Fig 1D, Fig S2D**). Limited proteinase-K digestion of LRH-1 ligand binding domain in the presence of bilirubin is delayed (**Fig 1C, Fig S3A**), also suggesting bilirubin binds LRH-1. Nuclear receptor ligands function to regulate transcription by inducing interaction with transcriptional coregulators^34,54,55^, we determined the bilirubin-induced apparent dissociation constants (*K_d_*) of LRH-1 for three established LRH-1 transcriptional coregulator peptides (SHP, PGC1α and DAX1) by fluorescence anisotropy, showing saturable binding curves (**Fig 1E**) and altered LRH-1 apparent *K_d_* relative to DMSO vehicle negative control (**Fig 1E**). These data suggest unconjugated bilirubin binds and regulates the function of the purified ligand-binding domain of LRH-1, *in vitro*.

**Figure 1.**
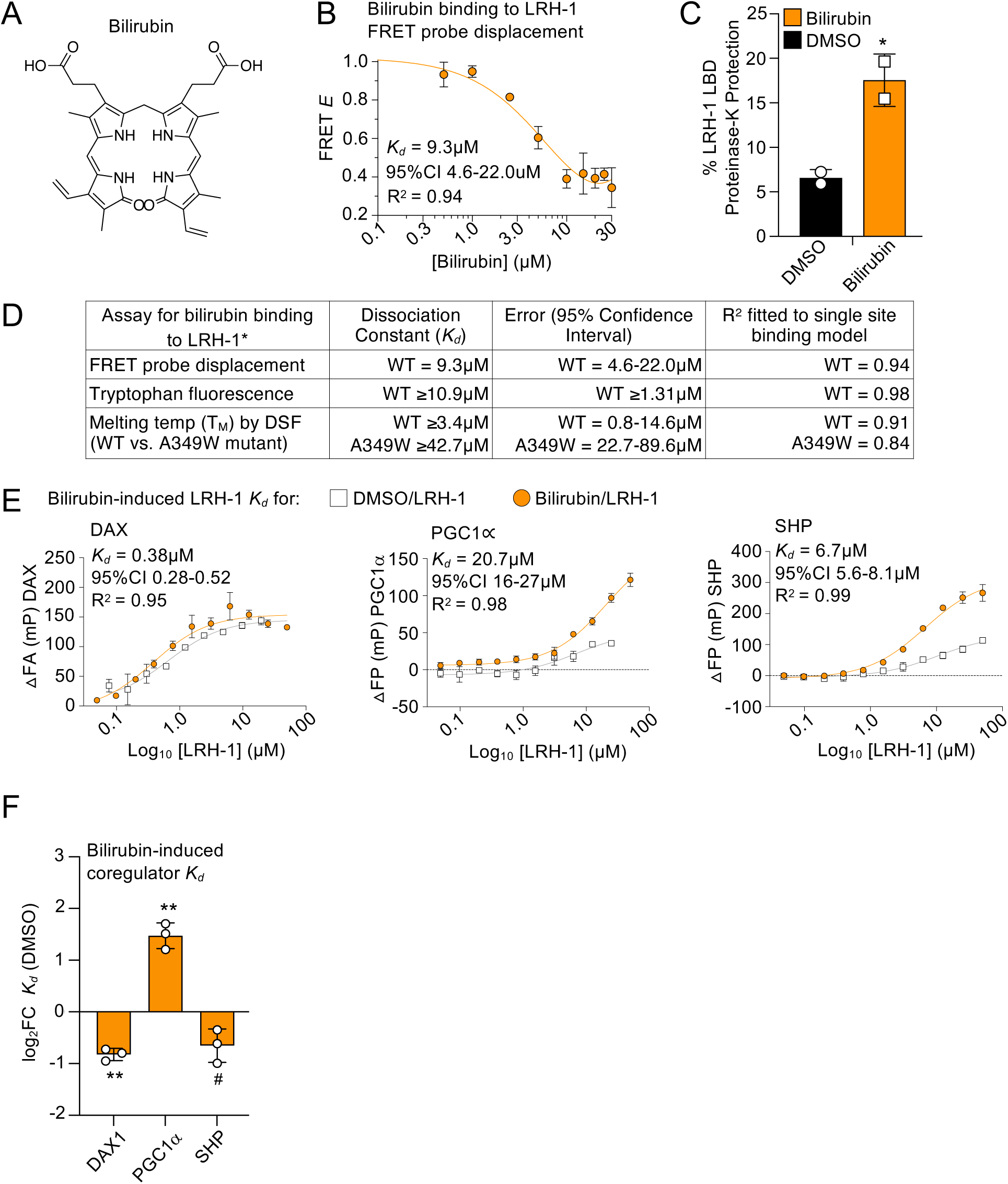
Bilirubin binds to LRH-1 ligand binding domain, regulating LRH-1 function *in vitro*. **A.** Chemical structure of unconjugated bilirubin. **B**. Phospholipid-probe displacement FRET assay, apparent *K_d_* of bilirubin for human LRH-1 ligand binding domain (LBD) is 9.3µM. **C.** Quantitation of LRH-1 LBD Proteinase-K protection assays in the presence of bilirubin or DMSO control, over a 60 min time course, bilirubin significantly protected LRH-1 in ANOVA across all time points (*p*=0.0458, 30 min shown here with full gel images in supplemental). **D**. Table of 3 independent assays used to determine bilirubin binding to LRH-1 by FRET displacement (9.3µM), internal tryptophan fluorescence (10.9µM) and melting temperature shift assay of indicated wild-type LRH-1 LBD (3.4µM) or the A349W pocket mutant (42.7µM). **E.** Change in fluorescent peptide anisotropy as a function of [LRH-1] to determine the indicated bilirubin-induced apparent *K_d_* of DAX1, PGC1α and SHP FAM-labeled peptides for LRH-1 LBD, **F**. also expressed as Log_2_ fold change (Log_2_FC) relative to DMSO control, bilirubin alone had no detectable effect on peptide anisotropy in the absence of LRH-1. In all panels \**p_adj_*<0.05, \*\**p_adj_*<0.01, in ordinary one-way ANOVA Dunnett’s corrected for multiple comparisons; ns is not significant; *^#^p<*0.05 by uncorrected t-test. These data suggest bilirubin binds the purified LRH-1 ligand binding domain and regulates the function of LRH-1 *in vitro*.

### Bilirubin stably interacts with LRH-1 *in silico*

Attempts to co-crystalize the bilirubin/LRH-1 complex have proven challenging, so we tested if bilirubin binding to the canonical ligand binding site in LRH-1was stable *in silico*. We used Rosetta to generate 2000 docked poses of the LRH-1/bilirubin complex (**Fig 2A**, **Spreadsheet 1**), the ten lowest-energy poses were used to calculate RMSDs against all poses (**Fig 2A**, **Fig S4-S5**), **Spreadsheet 2**), producing three groups of similar poses (**Fig 2B, Fig S4C-D**). Pose #1473 was the lowest energy pose in the most populated group (**Fig 2C**) and was used in 0.5µs molecular dynamics (MD) simulations (**Fig 2D-E**). Bilirubin interaction with LRH-1 was stable over the entire MD simulation, associating with changes to LRH-1 conformation (**Fig S5A-D, Video 1**). A second 0.5µs MD simulation starting with pose #1863 (**Fig 2B**) also showed stable interaction of bilirubin with LRH-1 over the entire simulation (**Video 2**). We next explored if changes to LRH-1 that occurred during the simulation in the presence of bilirubin might alter LRH-1 interactions with coregulators *in silico*, as 83.6% of MD frames clustered into a single state of LRH-1 (**Fig 2F**, **Table 1**). To explore this, the centroid of the most populated MD cluster was docked to DAX1, PGC1α and SHP peptides using Rosetta (**Fig S6**), suggesting bilirubin binding alters interaction of LRH-1 with all three coregulators *in silico* (**Fig 2G**). Similar results were obtained from the second MD simulation (#1873 as T_0_, **Table 2**), except PGC1α binding was not significantly altered (**Fig S7A-B**). In an orthogonal approach, we docked bilirubin to LRH-1 without specifying any ligand binding site *a priori* using PyRx, which also predicted bilirubin to bind the canonical ligand binding pocket of LRH-1 (**Fig S8A**). The blind-docked bilirubin was also stable over another independent 50ns MD simulation (**Fig S8B-E, Spreadsheet 3-4**) and the lowest energy state of LRH-1 from this simulation had altered interactions with PGC1α and SHP coregulator peptides *in silico* (**Fig S8F**). Negative-control docking runs of bilirubin to mouse Lrh-1 and the A349W mutant of human LRH-1 predicted bilirubin will not bind the canonical binding site in these variant LRH-1 proteins (**Fig S9A-B**), consistent with previous wet lab studies suggesting these LRH-1 variants bind ligand less, if at all^32,56,57^. These data suggest bilirubin binding LRH-1 is stable and can regulate LRH-1 functions *in silico*.

**Figure 2:**
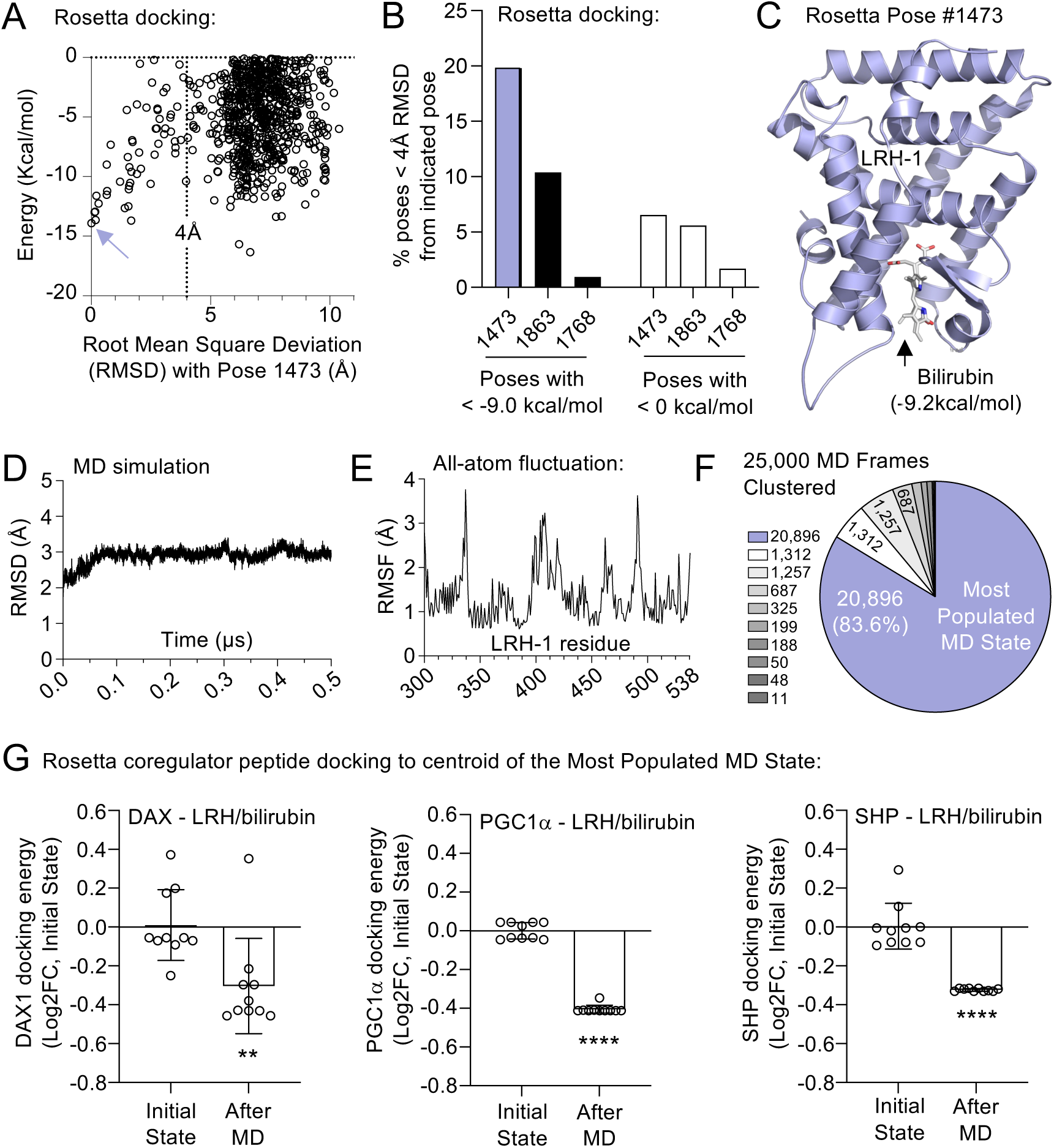
In silico analyses predict bilirubin stably interacts with the canonical ligand binding site in LRH-1. **A**. Calculated energy of all Rosetta-docked poses of bilirubin docked to LRH-1 with negative binding energy, plotted as a function of the root mean squared deviation of each structure from pose #1473 (blue arrow). Similar analyses were executed on a total of 10 low-energy poses (See supplemental). **B**. Percentage of all poses analyzed with RMSD< 4.0Å from indicated low-energy poses (#1473, #1863, #1768). **C**. Cartoon model of the Rosetta-consensus pose that was lowest energy in the most frequently populated group of poses. **E**. Following 0.5µs AMBER molecular dynamics (MD) simulation using pose #1473 as the initial state, calculated all-atom root mean square deviation (RMSD) is shown over the entire MD simulation. **F**. All-atom root mean square fluctuation (RMSF) of indicated LRH-1 amino acids over the entire MD simulation. **G**. Clustering of 25,000 states from the MD simulation, the centroid of the most populated state (frame 15783) represented 83.6% of all MD simulated states, used in panel H for peptide docking. **H**. Log2 fold change of the Rosetta energy score function of indicated coregulator peptides docked to LRH-1/bilirubin complex before (pose 1473, “Initial”) or after MD simulation (frame 15783, “MD”). Error is the standard deviation between 10 independent Rosetta peptide docking runs, these t-tests were Bonferroni-corrected for multiple comparisons as the docking runs were computed simultaneously (\*\**p_adj_*<0.01, \*\*\*\**p_adj_*< 0.0001). These data suggest the interaction of bilirubin with LRH-1 is stable *in silico*.

**Table 1.** Clustering output of 0.5µs MD frames, Rosetta pose 1473 used as T_0_. Bilirubin was docked to LRH-1 using Rosetta, and output pose 1463 was selected as time zero (T0) for 0.5µs MD simulation, detailed in methods. Here, 25,000 of the 50,000 total MD frames (every other frame) were clustered using a bottom-up hierarchical agglomerative approach with average linkage. The most populated cluster (1473-1) included 20,896 of the 25,000 frames, representing 83.6% of all analyzed frames. The mean distance between atoms in 1473-1 was 1.348Å, standard deviation was 0.190Å. Frame 15783 was the centroid of 1473-1, having the lowest cumulative distance to every other point in 1473-1, this frame was then used for coregulator peptide docking analyses. The average distance of 1473-1 to all 9 other clusters was 2.209Å. These data suggest frame 15783 from cluster 1473-1 is the most highly occupied state over the course of this 0.5µs MD simulation.

| Cluster | Frames in Cluster | Fraction of Frames | Mean distance (Å) | Standard Deviation ($\sigma$) | Centroid frame | Mean C-α distance (Å) |
| --- | --- | --- | --- | --- | --- | --- |
| 1473-1 | 20896 | 0.836 | 1.348 | 0.190 | 15783 | 2.209 |
| 1473-2 | 1312 | 0.052 | 1.308 | 0.230 | 1866 | 2.099 |
| 1473-3 | 1257 | 0.050 | 1.286 | 0.235 | 14805 | 2.416 |
| 1473-4 | 687 | 0.027 | 1.245 | 0.209 | 399 | 2.100 |
| 1473-5 | 352 | 0.014 | 1.100 | 0.157 | 2430 | 1.982 |
| 1473-6 | 199 | 0.008 | 0.000 | 0.000 | 20271 | 2.362 |
| 1473-7 | 188 | 0.008 | 1.266 | 0.219 | 895 | 2.030 |
| 1473-8 | 50 | 0.002 | 1.210 | 0.064 | 25 | 2.540 |
| 1473-9 | 48 | 0.002 | 1.184 | 0.000 | 850 | 2.334 |
| 1473-10 | 11 | 0.000 | 0.000 | 0.000 | 5 | 2.061 |

**Table 2.**
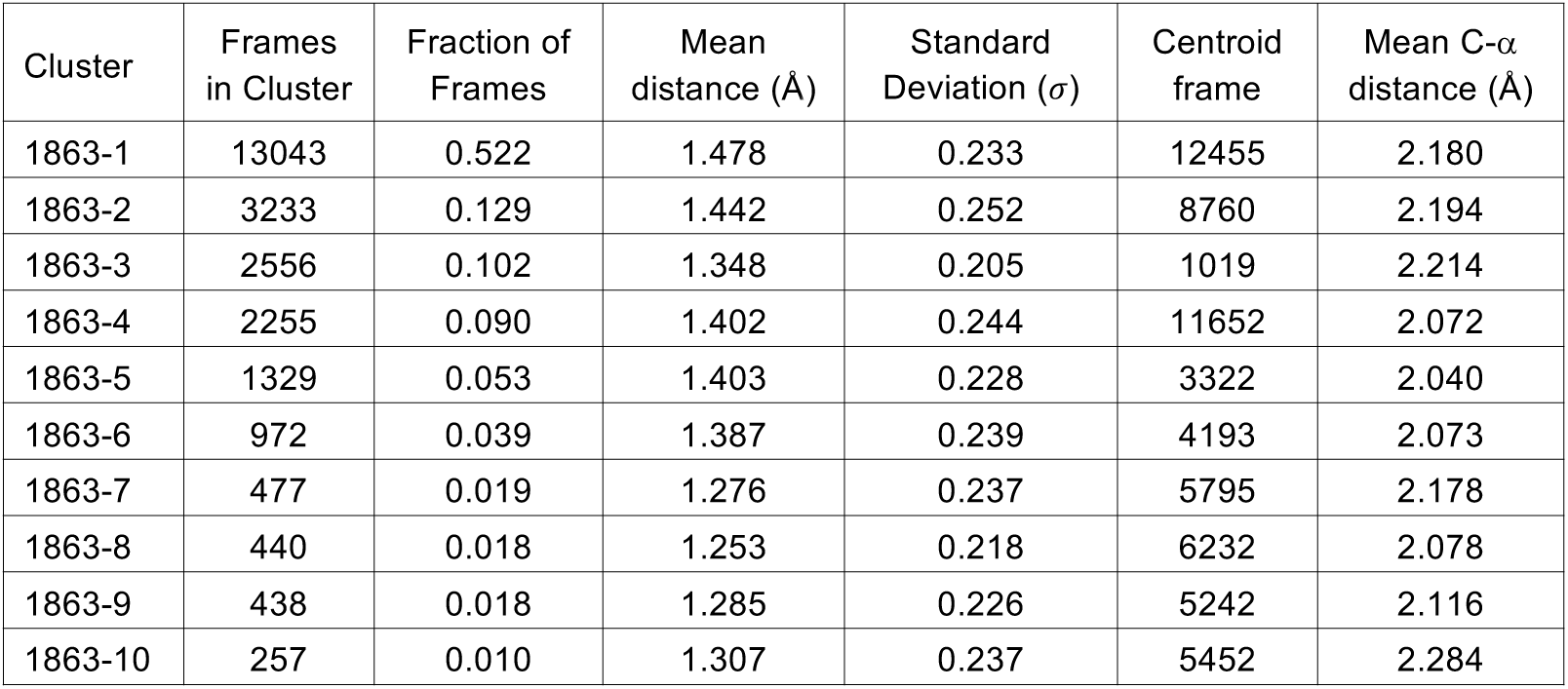
Clustering output of 0.5µs MD frames, Rosetta pose 1863 used as T_0_. Bilirubin was docked to LRH-1 using Rosetta as above in Table 1, but now output pose 1863 was selected as time zero (T0) rather than 1473, for an independent 0.5µs MD simulation. 25,000 of the 50,000 total MD frames were clustered using a bottom-up hierarchical agglomerative approach with average linkage. This analysis showed the top four clusters (1863-1, 1863-2, 1863-3 and 1863-4) all combined to represent 84.3% all frames, similar representation as the single 1473-1 cluster in Table 1 (83.6%). The centroid frame of 1863-1 was used for coregulator peptide docking analyses (12455). Thus, the MD output using 1863 as T0 for MD, was distributed across 4 distinct structural clusters of LRH-1 whereas the MD output using 1473 as T0 converged on a single conformation. Together, these clustering analyses suggest that frame 15783 from the 1463-1 cluster was the most populated single state derived from the MD simulations.

| Cluster | Frames in Cluster | Fraction of Frames | Mean distance (Å) | Standard Deviation ($\sigma$) | Centroid frame | Mean C-α distance (Å) |
| --- | --- | --- | --- | --- | --- | --- |
| 1863-1 | 13043 | 0.522 | 1.478 | 0.233 | 12455 | 2.180 |
| 1863-2 | 3233 | 0.129 | 1.442 | 0.252 | 8760 | 2.194 |
| 1863-3 | 2556 | 0.102 | 1.348 | 0.205 | 1019 | 2.214 |
| 1863-4 | 2255 | 0.090 | 1.402 | 0.244 | 11652 | 2.072 |
| 1863-5 | 1329 | 0.053 | 1.403 | 0.228 | 3322 | 2.040 |
| 1863-6 | 972 | 0.039 | 1.387 | 0.239 | 4193 | 2.073 |
| 1863-7 | 477 | 0.019 | 1.276 | 0.237 | 5795 | 2.178 |
| 1863-8 | 440 | 0.018 | 1.253 | 0.218 | 6232 | 2.078 |
| 1863-9 | 438 | 0.018 | 1.285 | 0.226 | 5242 | 2.116 |
| 1863-10 | 257 | 0.010 | 1.307 | 0.237 | 5452 | 2.284 |

### Bilirubin regulates a reporter specific for LRH-1, dependent on the bilirubin membrane transporter and a ligand-binding competent LRH-1

We next tested if bilirubin could induce LRH-1 activation of an LRH-1-specific *Cyp7A1* luciferase reporter in heterologous HEK293T cells. Previous studies showed cellular import of radiolabeled unconjugated bilirubin required ectopic expression of the bilirubin membrane transporter *SLCO1B1* in HEK cells^58,59^. Without *SLCO1B1*, 50µM bilirubin treatment for 24 hours did not change LRH-1 reporter activity (**Fig 3A**), but co-expression with *SLCO1B1* licensed bilirubin activation of the LRH-1 specific reporter (**Fig 3B**) suggesting bilirubin must be intracellular to activate LRH-1 in this assay. That the reporter required co-expression of LRH-1 suggests no factors endogenous to HEK293T cells are mediating activation of the luciferase reporter (**Fig 3A-B**). Further, the LRH-1 hypomorphic “pocket mutants” (A349F and A349W, **Fig 3C**) previously shown to prevent ligand binding to LRH-1^53^, were not regulated by bilirubin treatment even in the presence of *SLCO1B1* (**Fig 3B**). Under three optimized and independent transfection conditions, bilirubin activated the LRH-1-specific reporter in a concentration-dependent manner (**Fig 3D**), with a cellular EC_50_ of 28.9µM (95%CI = 9.5 -118.9µM). Since bilirubin does not activate the reporter in the absence of LRH-1 co-expression (**Fig 3A-B**), bilirubin induction of the reporter is dependent on LRH-1. These data suggest bilirubin activates an LRH-1-specific reporter in human HEK cells, dependent on the ability of LRH-1 to bind ligand and further dependent on co-expression with bilirubin transporter *SLCO1B1*.

**Figure 3:**
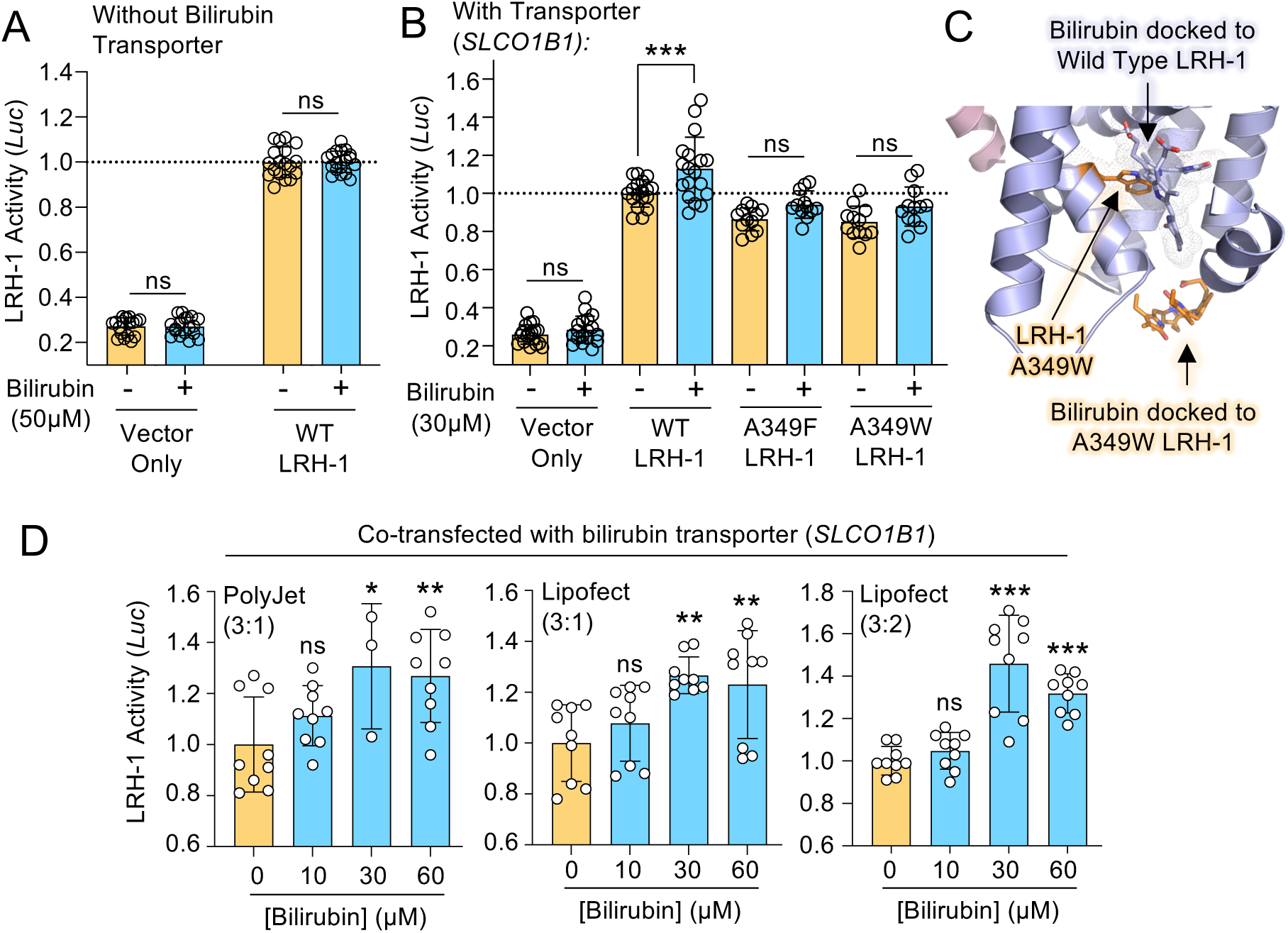
Bilirubin activates LRH-1 in living cells, dependent on co-expression with the plasma membrane bilirubin transporter SLCO1B1. **A**. *CYP7A1-*Lucifierase LRH-1 reporter assay in HEK293 cells transiently expressing all plasmids, in the absence of co-transfecting the bilirubin transporter *SLCO1B1.* Note bilirubin treatment has no effect on LRH-1 reporter activity despite presence at high concentration in the media. **B**. Same assay as in A., however a plasmid expressing SLCO1B1 has been added to the transfection, along with indicated LRH-1 wild type or LRH-1 pocket mutants (A349F and A349W), previously shown to bind ligand less than WT LRH-1. Regardless of the method of statistical analysis these LRH-1 mutants were not regulated by bilirubin. **C**. PyRx models of bilirubin docked to either WT LRH-1 (purple) or A349W mutant LRH-1 (orange), illustrating the predicted steric clash between mutant A349W residue and bilirubin. Bilirubin is not predicted to bind inside the canonical ligand binding pocket of A349W mutant LRH-1, the pink helix on left is the PGC1a peptide from PDB:6OQX. **D**. Concentration-dependent activation of LRH-1 by bilirubin, under 3 independent reporter transfection conditions, all in the presence of *SLCO1B1*.

### Bilirubin selectively regulates transcripts from endogenous LRH-1 ChIP-seq target genes in human HepG2 cells

While the luciferase system had utility testing bilirubin specificity for LRH-1, human HepG2 cells model endogenous LRH-1 regulation of endogenous target genes^20,60,61^. We generated HepG2 cells that stably express the SLCO1B1 bilirubin transporter, which sensitized the cells to bilirubin (50µM, 24 hours) induction of the *CYP7A1* target gene of LRH-1 (**Fig 4A**). Transcriptomes after bilirubin treatment (50µM, 24 hours) showed differential expression of 184 transcripts by RNA-seq (*p_adj_*>0.05 and Log_2_FC±1.0, **Fig 4B, Fig S10A-B, Spreadsheet 5**), and bilirubin regulated several known LRH-1 target genes by RT-qPCR (**Fig 4C**).

**Figure 4:**
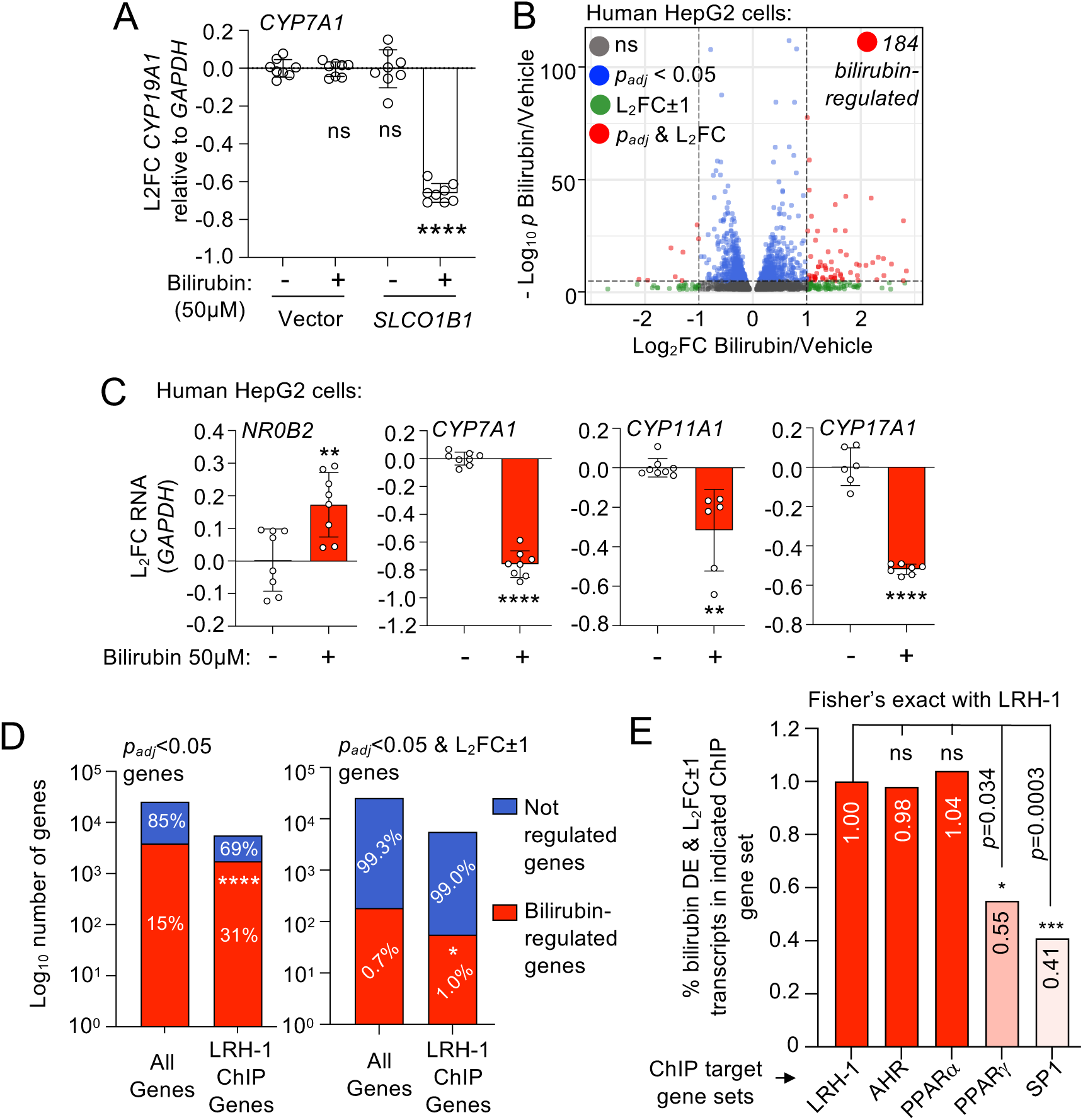
The bilirubin-regulated transcriptome in HepG2 cells is enriched in LRH-1 ChIP-seq target genes. **A**. RT-qPCR of RNA from HepG2 cells stably expressing either vector or *SLCO1B1*, treated with DMSO vehicle or 50µM bilirubin for 24 hours, showing the known LRH-1 target gene *CYP7A1* requires stable *SLCO1B1* expression for the bilirubin response. **B**. Volcano plot of the bilirubin-regulated transcriptome in HepG2 cells stably expressing the bilirubin membrane transporter *SLCO1B1*, treated with either vehicle or 50µM bilirubin for 24 hours; 184 transcripts are differentially expressed by bilirubin treatment (*p_ad_*_j_<0.05 and Log_2_ fold change (L_2_FC) at least ±1.0) **C.** RT-qPCR of established LRH-1 target genes from HepG2 cell RNA treated identically as in panel A (\*\**p*<0.01, \*\*\*\**p*<0.0001 by t-test). **D.** Contingency analysis comparing the number of all quantifiable genes in the transcriptome that are bilirubin regulated to the number of all quantifiable LRH-1 ChIP-seq target genes that are bilirubin regulated. Left panel shows distribution of the *p_adj_*<0.05 bilirubin-regulated genes, right panel shows distribution of the *p_adj_*<0.05 & Log_2_FC±1 bilirubin-regulated genes (\*\*\*\**p*<0.0001, \**p*=0.0343 by Fisher’s exact test), thus LRH-1 ChIP-seq target genes are enriched in bilirubin regulated genes when compared to all other genes in the quantifiable transcriptome. **E.** Comparing enrichment of bilirubin-regulated genes with known bilirubin nuclear receptors and other indicated ChIP-seq datasets, all ChIP data were from HepG2 cells. Enrichment on the left axis is the percentage of bilirubin-regulated transcripts (*p_ad_*_j_<0.05 & Log_2_FC±1) in each indicated ChIP gene set. Enrichment in the LRH-1 ChIP-seq dataset is compared individually to enrichment of bilirubin-regulated genes in each ChIP dataset for two positive control data sets (AHR and PPARα) and two negative control data sets (PPARγ or SP1). Several reports suggest bilirubin regulates AHR and PPARα, the above analysis suggests LRH-1, AHR and PPARα are all direct targets of bilirubin action in HepG2 cells. These data suggest the bilirubin-regulated transcriptome in HepG2 cells is enriched in direct LRH-1 ChIP-seq target genes.

Previous studies have identified direct LRH-1 target genes by chromatin immunoprecipitation (ChIP)-seq in HepG2 cells^20^(**Spreadsheet 6**), LRH-1 ChIP-seq gene sets were significantly enriched in bilirubin-regulated genes compared to all quantifiable genes in the transcriptome (**Fig 4D, Spreadsheet 7**), suggesting bilirubin selectively regulates transcripts from genetic loci that recruit LRH-1. However, HepG2 cells also express other known bilirubin receptors, specifically the aryl hydrocarbon receptor (AHR)^49,62^ and PPARα^50,51^, whose direct target genes also likely contribute to the bilirubin-regulated transcriptome. To compare enrichment of the bilirubin-regulated transcriptome in ChIP-target genes for all these bilirubin receptors, we used ChIP-datasets from the ENCODE^63^ project and other published sources^64^ all in HepG2 cells, to compare enrichment of bilirubin-regulated genes in the LRH-1^61^, AHR^64^ and PPARα^65^ ChIP data sets. All three of these ChIP gene sets were significantly enriched in bilirubin regulated target genes, with no significant differences in enrichment between LRH-1 *vs*. AHR or PPARα (**Fig 4E, Fig S11A, Spreadsheet 7**), suggesting these nuclear receptors all participate in the bilirubin transcriptional response^49,50^. Negative control ENCODE HepG2 ChIP-seq gene sets from the transcription factors PPARγ and SP1^63^ in HepG2 cells were significantly less enriched in bilirubin regulated genes (**Fig 4E, Fig S11B**), PPARγ and SP1 are not thought to bind bilirubin. To determine if serum in the media might affect induction of LRH-1 target genes, we confirmed that bilirubin activated the LRH-1 target gene *NR0B2* when these HepG2 cells were cultured in serum-free media, by RT-qPCR (**Fig S12**). Together, these data suggest bilirubin regulates transcripts emanating from direct LRH-1 target genes, as frequently as other more well-established bilirubin nuclear receptors.

### Genetic downregulation of several genes attenuates the bilirubin-regulated transcriptome in HepG2 cells

The above data suggest the bilirubin-regulated transcriptome is enriched in LRH-1 ChIP-seq target genes, we next asked if genetic downregulation of LRH-1 would attenuate bilirubin-induced gene expression. Transfection of siRNA against human LRH-1 (*NR5A2*) for 96 hours significantly downregulated expression of *NR5A2* transcripts in HepG2 cells (**Fig S13A**), as expected. Gene profiling in the presence of control siRNA (siCON) revealed 171 transcripts differentially expressed by bilirubin treatment (50µM for 24hrs, **Fig 5A, Fig S13B-C, Spreadsheet 8**), however in the presence of siRNA against LRH-1 (*NR5A2*), only 8 transcripts were differentially expressed by identical bilirubin treatment (**Fig 5B, Spreadsheet 9**). These data suggested that full bilirubin-induced gene expression requires full expression of LRH-1 in HepG2 cells, however as previously noted, HepG2 cells express other putative bilirubin receptors, and knockdown of PPARα in HepG2 cells also attenuated the response to bilirubin in a previous publication^66^. We tested the effect of siRNA against PPARα, as well as negative controls PPARγ and SP1. Surprisingly, all of these siRNAs attenuated the bilirubin-induced transcriptome in these HepG2 cells similar to siRNA against LRH-1 (**Fig 5C-E**). Bilirubin was able to regulate the transcriptome in the presence of *siNR3C2* treatment (**Fig 5F**), suggesting the attenuation of bilirubin response was not due to transfection of any siRNA sequence, also supported by the *siCON* control (**Fig 5A**). It is unclear why *siNR3C2* (Mineralocorticoid Receptor) enhances the bilirubin response. Analyses of the transcriptomes revealed that in all conditions where the bilirubin response was attenuated, LRH-1 transcripts had decreased (**Fig 5G-J**), including the condition of siRNA against PPARα, suggesting PPARα and LRH-1 might co-regulate each other within a gene network. Indeed, recent data suggest PPARα and LRH-1 are in a tightly-coregulated network in the livers of human patients, through regulation of enhancer sequences^67^. Thus, although loss of either LRH-1 or PPARα prevented bilirubin regulation of the HepG2 transcriptome, disentangling any specific LRH-1 and PPARα responses may be difficult to achieve using genetic knockdown models.

**Figure 5:**
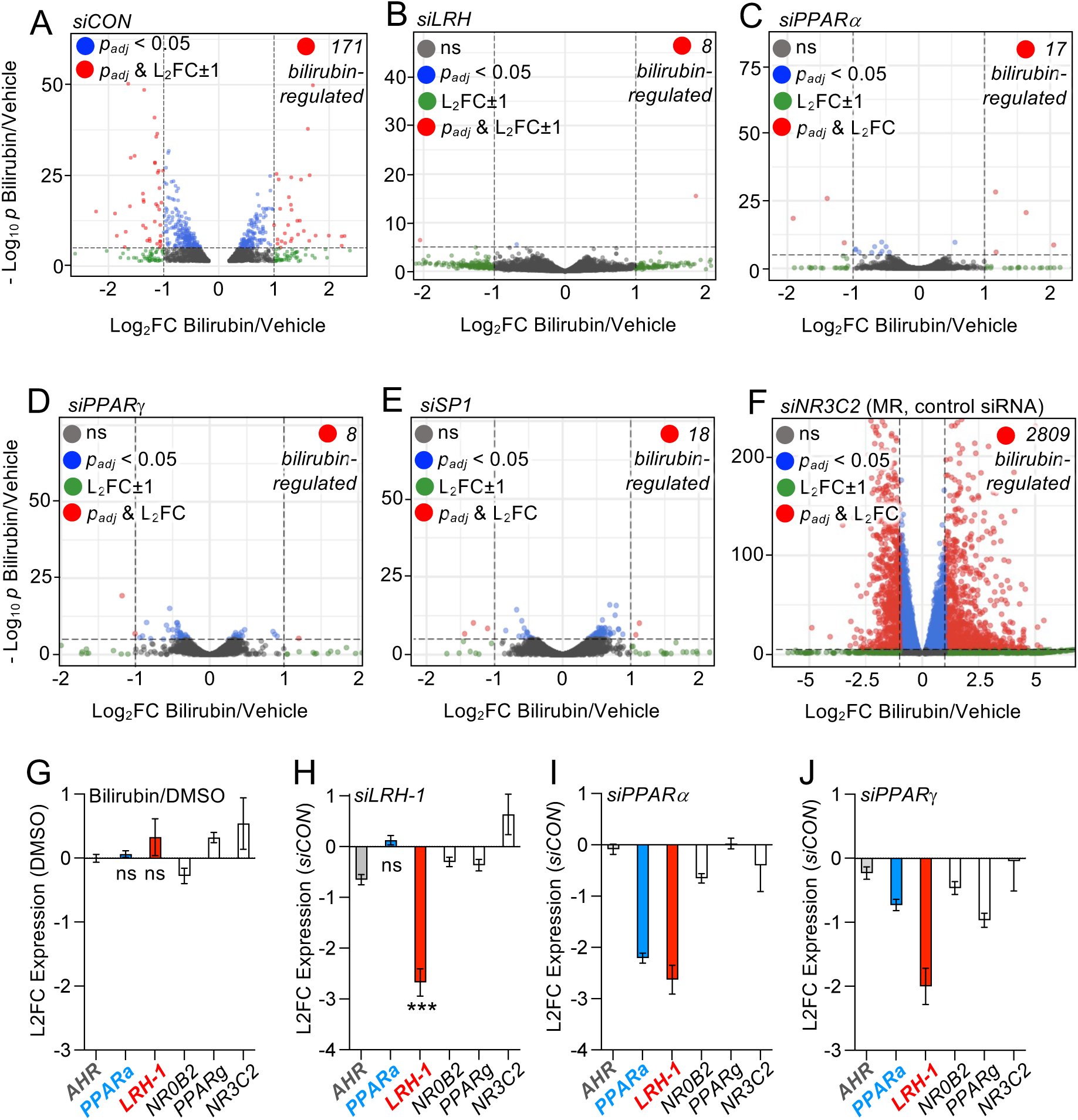
Knockdown of PPARa or PPARg also downregulates LRH-1, in HepG2 cells. **A.** Volcano plot of transcripts induced by 24h, 30µM bilirubin**. B-F** Volcano plots of 24h, 30µM bilirubin-induced transcriptomes from HepG2 pre-treated for 72 hours with indicated siRNAs, showing the bilirubin-induced transcriptomes are attenuated in all conditions except panel F (siRNA against *NR3C2*, MR). **G**. Log2 fold changes of relevant individual transcripts from the panel A transcriptome, induced by 24h, 30µM bilirubin**. H-J.** Log2 fold changes of indicated transcripts regulated by indicated siRNA labeled on top of each graph, relative to control siRNA (siRNA / siCON), showing that in all conditions where LRH-1 is downregulated, the bilirubin-induced transcriptomes were attenuated (top panels), *\*\*\*p_adj_*<0.001 between *LRH-1* and *PPARα* t-test, Dunnett’s corrected. These data suggest siRNA knockdown of PPARa or PPARg results in concomitant knockdown of LRH-1 transcripts, suggesting genetic approaches understanding bilirubin mechanism of action in HepG2 cells are insufficient..

### A chemical competitor of LRH-1 attenuates the bilirubin-regulated transcriptome in HepG2 cells

PPARα and LRH-1 appear to be genetically co-regulated in an integrated gene expression network^67^ (**Fig 5**). Further, any genetic experiment that removes LRH-1 only address if LRH-1 expression is required for the bilirubin response, not if bilirubin binding to LRH-1 is required for the transcriptional response. To begin to address these points, we tested if the LRH-1 chemical competitor RJW100^68^ could attenuate bilirubin regulation of the HepG2 transcriptome. A previously published crystal structure of the RJW100-LRH-1 complex demonstrates RJW100 binds the canonical ligand binding site in LRH-1^69^ and displaces ligands bound to this site in LRH-1^53,69^ (**Fig 6A**). A two-fold molar excess of RJW100 (100µM) attenuated 50µM bilirubin-induced transcriptome in HepG2 cells after 24 hours co-treatment of both these ligands (**Fig 6B-C, Spreadsheet 10**), with only 6 transcripts differentially expressed in response to bilirubin in the presence of RJW100 (*p_adj_*>0.05 and Log_2_FC±1.0, **Fig 6C, Spreadsheet 11**). These data suggest the transcriptional effects of 50µM bilirubin are attenuated by 100µM RJW100, a small molecule that directly binds the canonical ligand binding site in LRH-1. Together, the data suggest bilirubin behaves as a regulatory ligand for LRH-1.

**Figure 6:**
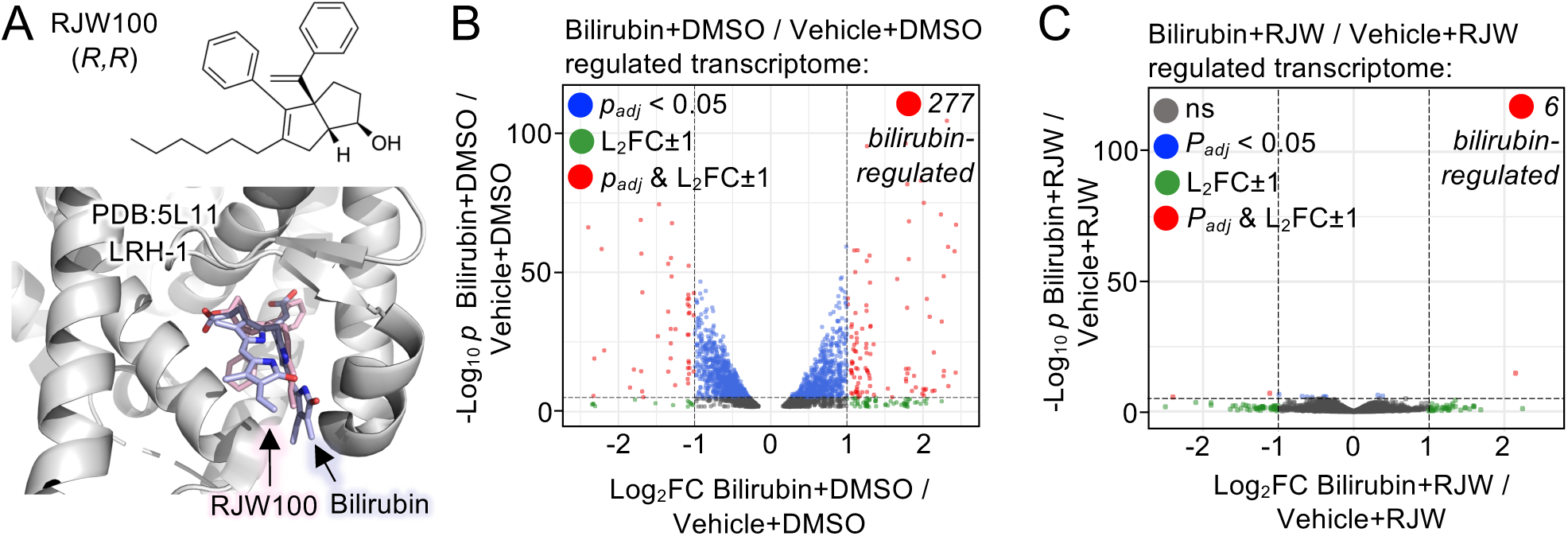
A chemical competitor of LRH-1 attenuates bilirubin regulation of the HepG2 transcriptome. **A**. Published crystal structure of LRH-1 bound to the small molecule RJW100 (RJW, PDB:5L11, gray), which is superposed with the computationally-docked position of bilirubin, predicting strong steric clashes between RJW100 (pink sticks) and bilirubin (blue sticks). Previous studies demonstrated RJW100 displaces ligands from the ligand binding pocket of LRH-1. **B**. Volcano plot of the 24 hour, 50uM bilirubin-regulated transcriptome of HepG2 cells stably expressing the SLCO1B1 bilirubin membrane transporter, in the absence or **C.** presence of 2-fold molar excess RJW100 competitor (100uM), “Vehicle” indicates bilirubin vehicle (100uM beta-cyclodextrin in DMSO), whereas “DMSO” alone is the vehicle for RJW100. These data suggest that a chemical competitor of the LRH-1 ligand-binding site attenuates bilirubin-regulation of the HepG2 transcriptome.

## Discussion

Here we present multiple, orthogonal lines of evidence suggesting bilirubin regulates LRH-1 functions *in vitro*, *in silico* and in human cells. That genetic downregulation of LRH-1 almost completely attenuated all transcriptional response to bilirubin in HepG2 cells (**Fig 4A-C**), coupled with similar effects seen with a chemical competitor of LRH-1 (**Fig 6B-C**), provide the first functional links between LRH-1 and bilirubin-induced transcriptional responses. A limitation of this study is our data suggesting that PPARα knockdown also downregulates LRH-1 in HepG2 cells, making it difficult to disentangle the effects of bilirubin on one receptor or another challenging, however that an LRH-1 chemical competitor also interferes with bilirubin activation of LRH-1 target genes, suggests LRH-1 acts as a receptor for bilirubin.

It is tempting to speculate on teleological reasons bilirubin might regulate LRH-1, given the roles both bilirubin and LRH-1 have in liver physiology. The simplest hypothesis is that LRH-1 senses bilirubin to help protect from high levels of bilirubin accumulation in hepatocytes. Phenotypes connecting bilirubin to LRH-1 in mice have not been reported, however mouse Lrh-1 binds ligands poorly if at all^56^, suggesting other ligand-sensing mechanisms likely exist in mice that are independent of mouse Lrh-1.

Gene set enrichment analyses suggest bilirubin represses cholesterol biosynthetic genes and activates genes associated with lipid efflux (**Fig S13, Spreadsheet 12**) consistent with shared roles for LRH-1 in known bilirubin-response pathways^19,47,61,70^. However, establishing those more physiological connections in the liver is outside the scope here. Bilirubin is a component of bile when conjugated to glucuronic acid (conjugated bilirubin) in hepatocytes, while unconjugated bilirubin serves as the metabolic precursor^44,71^.

LRH-1 has important roles in hepatic bile acid metabolism, activation of bile acid anabolic genes is regulated by a pathway involving LRH-1, Liver X Receptor (LXR) and Farnesoid X Receptor (FXR)^72,73^. The lack of a commercial source of conjugated bilirubin diglucuronide makes testing binding to LRH-1 difficult in the wet lab, however docking studies suggest unconjugated bilirubin binds LRH-1 with better affinity than conjugated bilirubin (**Fig S15**). We explicitly acknowledge that we have not tested all heme metabolites for binding to LRH-1, and LRH-1 may bind directly to conjugated bilirubin and/or biliverdin, which could obviously result in a very different influence on LRH-1 functions, which could provide clues as to the physiological consequences of bilirubin regulation of LRH-1.

We observed that bilirubin decreased expression of the direct LRH-1 target gene *CYP7A1* and other cholesterol-based bile acid biosynthetic genes (**Fig S14**). Speculating on why bilirubin might regulate LRH-1, LRH-1 might detect high ratios of unconjugated to conjugated bilirubin in hepatocytes, inducing the decreased expression of cholesterol metabolic enzymes to decrease the overall intrahepatic lipid load under the conditions of high levels of intrahepatic bilirubin. While bile acid feedback is well-known to occur independently of Lrh-1 in mice^74^, mouse Lrh-1 binds ligands poorly if at all^56^, and our docking studies show mouse Lrh-1 does not bind bilirubin *in silico* (**Fig S13**), together suggesting other genes are likely to drive bilirubin-sensing functions in rodents. The nuclear receptor CAR (*NR1I3*) has a clear role in mediating hepatic transcriptional responses to bilirubin in mouse models^75,76^ and CAR-specific genes are enriched in our human transcriptomes presented here (**Fig S14**), however up to 100µM bilirubin was unable to activate a CAR-specific luciferase reporter, suggesting bilirubin does not directly bind CAR^75^. Previous studies have shown bilirubin can activate an AHR-specific luciferase assay^49^. Bilirubin is also known to scavenge oxidative molecules^77^ and LRH-1 helps coordinate the redox state of the liver^78^ by controlling expression of redox-associated genes *Gpt2*, *Gls2* and *Got1*^79^, consistent with a potential role for LRH-1 sensing bilirubin to help properly modulate hepatocyte redox state. Although intriguing speculation, the physiological studies required to interrogate these hypotheses are beyond the scope of this manuscript.

Here, we identified bilirubin to regulate purified LRH-1 ligand binding domain functions *in vitro* (**Fig 1**), which is predicted to stably bind the canonical, orthosteric site in LRH-1 *in silico* (**Fig 2**). Bilirubin specifically regulated a reporter for LRH-1 in living human cells (**Fig 3**), as well as transcripts from endogenous LRH-1 target genes in HepG2 cells (**Fig 4**), which was dramatically attenuated by genetic downregulation of LRH-1 (**Fig 5**) or a chemical competitor of LRH-1 (**Fig 6**). Although these studies cannot directly address the relative contributions of LRH-1 and PPARα, our data do suggest bilirubin binds and regulates LRH-1 in addition to PPARα. Further studies will be required to determine the precise mode of bilirubin interaction with LRH-1, the structural mechanism of LRH-1 regulation and the physiological consequences of bilirubin regulation of LRH-1.

## Limitations of this study

With no crystal structure of the bilirubin/LRH-1 complex, atomic-resolution details of how LRH-1 is regulated by bilirubin remain unknown, although our data suggest bilirubin binds the purified ligand binding domain of LRH-1 and regulates *in vitro* functions of LRH-1. It is further unknown if bilirubin associates with LRH-1 purified from human tissues as these tissues are difficult to acquire and antibodies do not effectively purify LRH-1. Genetic studies in mice are unlikely to demonstrate physiological relevance because mouse LRH-1 does not bind ligand, suggesting other mechanisms exist in rodents to sense bilirubin levels.

## Methods

### Materials

Unconjugated bilirubin was purchased from ThermoFisher (catalog #A17522), resuspended in the dark in either DMSO for *in vitro* experiments using pure proteins, or 100uM beta-cyclodextrin in DMSO for all cell-based experiments. Chemicals were purchased from Fisher, Sigma, Avanti Phospholipids, Cayman Chemicals, Qiagen or Promega. pRSF2 full-length human LRH-1 (NCBI accession number NM_205860.3) construct was a gift from Robert Fletterick (University of California at San Francisco). Full-length human LRH-1 CMV mammalian expression vector was purchased from VectorBuilder (Chicago, IL). HepG2 and HEK293T cells were purchased from ATCC (HB-8065, CRL-3216). Plasmids expressing cDNAs for *SLCO1B1* and vector only control were purchased from VectorBuilder (Chicago, IL). RJW100 small molecule was custom synthesized by the Vanderbilt Institute for Chemical Biology according to the published synthesis^53^. *CYP7A1* firefly luciferase reporter construct was a gift from Holly Ingraham (University of California at San Francisco). *CYP7A1* promoter DNA oligonucleotide was purchased from IDT (Coralville, IA). FAM-labeled coregulator peptides were purchased from New England Peptide (Gardner, MA).

### LRH-1 ligand binding domain (LBD) protein expression and purification

A wild type human LRH-1 (*NR5A2*) ligand-binding domain (LBD) plasmid construct (residues 266-541, UniProt ID O00482) was cloned into isopropyl 1-thio-b-galactopyranoside (IPTG)-inducible pET vector with an N-terminal His-tag and expressed in *E. coli* BL21 dE3 using standard methods. Briefly, *E. coli* BL21 dE3 were transformed with an ampicillin-resistant LRH-1 LBD expression plasmid, grown at 37°C until OD_600_ = 0.8 and induced with 1mM IPTG overnight at 15°C at 100RPM shaking. Cells were collected by centrifugation at 3000xg at 4°C for 20 minutes, the pellet resuspended in lysis buffer (20mM Tris (7.5), 150mM NaCl, 2mM CHAPS, 1X Peirce/Roche protease inhibitors), transferred to a 50mL conical and stored at -80°C. Cell pellets were thawed on ice, lysed by probe sonication and LRH-1 purified from the debris-cleared lysate using TALON metal affinity chromatography, followed by TEV cleavage of the 6X-His tag. This protocol regularly yields LRH-1 LBD protein that is 95% pure by SDS-PAGE, and readily crystalizes in the absence of added ligand. All constructs are available on request.

### LRH-1 *K_d_* determination for unconjugated bilirubin by FRET

A previously established FRET-based phospholipid displacement assay^80^was used to approximate the apparent *K_d_* of unconjugated bilirubin for human LRH-1 LBD. Briefly, purified wild-type LRH-1 LBD was labeled with Alexa Fluor-488 C5-maleimide (Life Technologies, Carlsbad, CA) by incubation of the protein with a 1:10 molar excess of the fluorophore label overnight at 4°C. Labeled protein was obtained as detected by electrophoresis, and unincorporated label removed by desalting until no excess unincorporated fluorophore could be detected as retained on desalting column by fluorescence intensity measurements. Dioleoyl phosphatidyl-ethanolamine (18:1,18:1) labeled on the headgroup with Rhodamine (#810150, Avanti Polar Lipids, Birmingham, AL) was used as a FRET acceptor ligand while AlexaFlour-488 on the LRH-1 LBD was the FRET donor, FRET efficiency (E) for this FRET pair was calculated using the equation, E = 1-(F_DA_/F_D_), where F_DA_ is fluorescence intensity of the donor (LRH-1 LBD) in the presence of acceptor (Rh-PE phospholipid probe) and F_D_ is fluorescence intensity of the donor in the absence of the acceptor upon donor excitation. The FRET signal was undetectable in the absence of LRH-1 protein. Fluorescence intensity of donor at λ_emm_=530 nm was measured following excitation at λ_exc_=480 nm. Unconjugated bilirubin was resuspended in DMSO to 10mM and serially diluted in DMSO, assay plates were prepared by transferring 60nL of the bilirubin stocks from source plates to a 384-well black non-binding plate using an Echo 550 acoustic dispenser, with the first and last two columns of each plate loaded with 60nL of DMSO vehicle as no-compound controls. 40uL of assay buffer (25mM Tris-HCl, pH 7.5) was then added onto the assay plate using Multidrop Combi Reagent Dispenser. 10uL of LRH-1-488 was then added using Agilent Bravo Liquid Handler and the plates incubated at room temperature overnight, the final [LRH-1-488] was 50nM. Using an Agilent Bravo Liquid Handler, 10uL of either Rh-PE stock or assay buffer were added to wells containing DMSO control wells and 10uL of RH-PE to wells containing various concentrations of unconjugated bilirubin, to a final concentration of 50nM LRH-1-488 donor and 50nM of (18:1) Rh-PE acceptor. Plates were then incubated at room temperature for an hour to equilibrate, spun at 1000xg for 5 mins and fluorescence intensities measured at λ_emm_ = 520nm following excitation at λ_exc_ = 480nm using a Biotek Neo multimodal plate reader. The FRET efficiency E was calculated using equation (1) above, where F_D_ is the fluorescence intensity of the donor-only sample, which is decreased to F_DA_ by the acceptor in the donor-acceptor sample. Data were fit to a one-site binding model (non-linear regression) in GraphPad Prism (GraphPad Software, Inc. La Jolla, CA) to determine the apparent *K_d_* and 95% confidence interval values. All raw FRET data are available on request.

### LRH-1 *K_d_* determination by melting temperature

Bilirubin alone had no effect on this assay, explained below. Wild-type or the “pocket mutant” (A349W) human LRH-1 LBD were incubated overnight at room temperature with 2-fold serially diluted bilirubin in buffer or a DMSO vehicle control, LRH-1 LBD was at 0.166 mg/mL (50nM) in 20mM Tris (8.0) with 1x SYPRO Orange (Invitrogen). Differential scanning fluorimetry melting curves were measured in 96-well plates using a CFX96 Touch RT-PCR System or a CFX Opus 96 RT-PCR System (Bio-Rad), all samples were heated over a temperature gradient from 15 °C to 95 °C at increments of 0.5 °C every 30 seconds, with a fluorescence scan at each temperature increment, for each of quadruplicate samples. Bilirubin alone (in the absence of LRH-1 protein but identically treated in all other respects) in these assays run in parallel had no signal above background throughout the entire melting curve. Raw data for LRH-1 were background subtracted and analyzed using the SimpleDSF viewer MATLAB script to determine the melting temperature (T_Melt_) of LRH-1 at each bilirubin concentration. The T_Melt_ values were then fitted to a sigmoidal dose response (variable slope) equation using GraphPad Prism to yield the *K_d_*.

### Protease protection assay

Purified human LRH-1 LBD (13.4uM) in 20 mM Tris-HCl, pH 7.5, 100 mM NaCl was incubated with either a 20-fold molar excess of bilirubin or an equal volume of DMSO vehicle alone overnight at room temperature. Following overnight binding, Proteinase K (7.5 ng/reaction) or an equal volume of buffer alone was then added to each mixture followed by incubation at 25°C for the indicated time points, aliquots were taken at 10, 30 and 60min during proteolysis and digestion stopped by addition of SDS-PAGE buffer and boiling at 95°C for 5 mins, followed by 10% SDS-PAGE and Coomassie staining of duplicate digestion reactions for each condition. Gel band intensities were quantified using Image J (NIH), ANOVA of the full-length, undigested band showed a significant difference between DMSO *vs*. bilirubin treated LRH-1 across all time points (*p*=0.0458), statistical analyses done in GraphPad Prism.

### Fluorescence anisotropy LRH-1 *K_d_* determination for coregulator peptides induced by bilirubin

Effects of bilirubin on the recruitment of transcriptional coregulator peptides to LRH-1 LBD were determined by fluorescence polarization/anisotropy following a similar protocol published method by Mays, et. al.^80^. Briefly, LRH-1 LBD was incubated with bilirubin in a 1:10 molar excess of bilirubin or an equal volume of DMSO vehicle overnight at 4°C. The complex was then transferred into 384-well black-walled plates and serially diluted in assay buffer (20-mM Tris-HCl, pH 7.5, 100mM NaCl) with 50nM FAM-labeled coactivator peptides, fluorescence polarization/anisotropy in each well measured using a Biotek Neo multi-modal plate reader (Biotek Instruments, Winooski, VT). The sequences of N-terminally FAM-labelled peptides used were: DAX1-3, 5Fam-QGSILYSLLMSAKQ-OH; PGC1α, 5Fam-EEPSLLKKLLLAPA-OH; and SHP-1, 5Fam-RPTILYALLSPSPR-OH and were all chemically synthesized (New England Peptide, Gardner, MA). Bilirubin had no detectable effects on fluorescence polarization/anisotropy in the absence of LRH-1 protein. All assays were conducted in triplicate. Data were fit to a single-site equilibrium binding equation using GraphPad Prism (GraphPad Software, Inc. La Jolla, CA) to determine the apparent *K_d_* and *B*_max_. All raw anisotropy data are available on request.

### RosettaLigand bilirubin computational docking to LRH-1

A crystal structure of human LRH-1 LBD (PDB:1YOK) was used for computational docking with bilirubin using RosettaLigand^81^. Prior to docking, bilirubin ligand preparation produced 200 energetically accessible conformations of bilirubin using conformer generator BCL::Conf^82^. RosettaLigand docking was performed to generate 2000 docked bilirubin-LRH-1 complexes. Of these 2000 poses two were selected (Pose #1473 and Pose #1863) for 0.5µs (500ns) MD simulation analyses, these two were selected based on the low Rosetta energy score functions (energy) and the frequency with which similar poses occurred among the 2000 output poses. To calculate frequency of each pose, the root mean square deviation (RMSD) of each pose was calculated in RosettaLigand and compared against the lowest 10 Rosetta energy score function poses as reference, all poses within 2Å RMSD from each other were grouped. Pose #1473 was the centroid of the most frequently populated group, pose #1863 was the centroid of the second most populated group, the third group of poses was eliminated as it was not comparably populated. All docked coordinates, binding energies and other output from RosettaLigand are available on request.

### 0.5µs MD Simulations

We performed a 0.5µs molecular dynamics simulations on the complex of LRH-1 bound to bilirubin created by Rosetta docking above. Complexes were prepped for MD simulation using the CHARMM-GUI^83^. Briefly, the complex was solvated in an octahedral box of explicit TIP3P water with a 10-Å buffer around the protein complex. 50 Potassium (K+) and 42 Cl-ions were added to neutralize the system’s charge and simulate physiologic conditions. SHAKE^84^ bond length constraints were applied to all bonds involving hydrogen. Nonbonded interactions were evaluated with a 9Å cutoff and calculated with a particle-mesh Ewald summation method^85^ ^86^. The MD system was first minimized for 100 steps using steepest descent followed by 5,000 steps of conjugate gradient minimization. The system was then heated to 300K over 125ps in the NVT ensemble using a step size of 1fs. After switching to the NPT ensemble, positional restraints on protein and ligand were gradually removed while Langevin dynamics with a collision frequency of 1.0ps**^-^**^1^ was used for temperature regulation. Finally, production MD was conducted for 0.5µs using a step size of 4 fs, constant pressure periodic boundary conditions, isotropic pressure scaling and Langevin dynamics. MD trajectories printed every 20ps and 25,000 frames were created for further analysis. All minimizations and 0.5µs MD simulations were performed with Amber17^87^ employing the ff14SB, TIP3P, GAFF2 force field for proteins, ligand and water molecules, respectively^88^. Analyses of the MD trajectories were performed using the CPPTRAJ module^89^ of AmberTools. Root mean square deviations (RMSD) were calculated for all non-hydrogen atoms of protein residues for each frame in the trajectory using the initial structure as the reference. Root mean square fluctuations (RMSF) were calculated on all non-hydrogen atoms of protein residues for each frame in the trajectory using the initial structure as the reference. All MD simulation output frame coordinates and energies are available on request.

### Cluster analysis of 25,000 MD simulated frames

To determine the most populated frames in each MD run, 25,000 frames from the 0.5µs MD simulations, the program CPPTRAJ module^89^ of AmberTools was used with hierarchical agglomerative and average-linkage clustering, clustering was completed when the minimum distance between clusters was greater than 3Å, and only 10 clusters remained. The cluster containing the most structures was the most populated cluster of MD frames in each run, and the single centroid structure output from AmberTools with the lowest distance to every other point in all structures within the cluster (Tables 1 and 2) was used for further Rosetta peptide docking below.

### Rosetta peptide docking to 0.5µs MD simulated states

LRH-1-coregulator interface analysis was performed for the above selected 0.5µs MD simulation frames using RosettaScripts^90^. Interfacial energy of the LRH-1-coregulator peptide was calculated for the most populated MD frame from the 0.5µs simulation and compared with the initial frame from the MD run. The interface analysis was performed on the complex of the selected MD frames with each of the three coregulator peptides; DAX1, SHP, and PGC1α. The template structures were relaxed using RosettaScripts to remove clashes identified by the score function as present in the crystal structure. Coregulator peptides were then oriented into an interaction orientation at the AF-2 (Helix 12) of LRH-1 that has been established crystallographically and minimization performed using RosettaScripts. Manual pre-orienting the coregulator-peptide into the established AF-2 region introduces investigator bias, however this dramatically decreased computation time by eliminating the coarse and fine resolution search stages of the docking but retaining the minimization stage. The protocol used here is otherwise identical to standard protein-protein docking that has been used in other studies^91–94^. This procedure generated ten models for each LRH-1/coregulator peptide, InterfaceAnalyzer mover in RosettaScripts calculated binding energy (ΔG kcal/mol) of each LRH-1-coregulator interface (ΔG_Separated) by separating and repacking the chains. This metric represents the change in Rosetta energy when the interfacial residues are separated compared to the energy when the same residues are complexed. All docked position coordinates and binding energies are available on request.

### Blinded/Global PyRx Docking

Unconjugated bilirubin was docked to the human LRH-1 LBD crystal structure (PDB: 1YOK) using PyRx^95^ in a blind docking approach. Before docking, bilirubin was energy minimized by 200 steps using a universal force field (UFF). The size of the grid box was set to X:50Å, Y:62Å, Z:47Å which encompassed the entire ligand binding domain of the LRH-1 protein, such that bilirubin had access to all potential binding sites in LRH-1 that are not the canonical ligand binding site, all PyRx docking data are available on request.

### Local PyRx docking and 50ns MD simulations

Unconjugated and conjugated bilirubin (Supplemental Figure S13) were docked into the human LRH-1 LBD crystal structure (PDB: 6OQX), or the same crystal structure containing the pocket mutant LRH-1 (A349W, Supplemental Figure S11A), or the mouse Lrh-1 LBD crystal structure (PDB:1PK5, Supplemental Figure S11B), using PyRx^95^. Before docking, bilirubin was energy minimized by 200 steps using a universal force field (UFF). The size of the grid box was X:22.7Å, Y:25.2Å, Z:26.1Å, centered on the canonical ligand binding pocket of LRH-1, all compounds and coregulator peptides that had been co-crystalized in the structures were removed prior to docking. The unconjugated bilirubin LRH-1 complex was then prepped for 50ns MD simulation using the CHARMM-GUI^96^. Briefly, the complex was solvated in an octahedral box of explicit TIP3P water with a 10-Å buffer around the protein complex. 49 Potassium (K^+^) and 41 Cl^-^ ions were added to neutralize the protein and simulate physiologic conditions.

SHAKE^84^ bond length constraints were applied to all bonds involving hydrogen. Nonbonded interactions were evaluated with a 9Å cutoff and calculated with a particle-mesh Ewald summation method^85,86^. The MD system was first minimized for 100 steps using steepest descent followed by 5,000 steps of conjugate gradient minimization. The system was then heated to 300K over 125ps in the NVT ensemble using a step size of 1fs. After switching to the NPT ensemble, positional restraints on protein and ligand were gradually removed while Langevin dynamics with a collision frequency of 1.0ps**^-^**^1^ was used for temperature regulation. Production MD was conducted for 50ns using a step size of 2fs, constant pressure periodic boundary conditions, isotropic pressure scaling and Langevin dynamics. All minimizations and 50ns MD simulations were performed with Amber17^97^ employing the ff14SB, TIP3P, GAFF2 force field for proteins, ligand and water molecules, respectively^98^. Analysis of the MD simulations was performed on 5000 frames using the CPPTRAJ module^89^ of AmberTools. RMSDs were calculated for all non-hydrogen atoms of protein residues for each frame in the trajectory using the initial structure as the reference, RMSF were calculated on backbone and all non-hydrogen atoms of protein residues for each frame in the trajectory using the initial structure as the reference. Out of 5000 output frames, frames #2635 and #4719 were selected as the lowest energy states and compared with the initial frame of the MD simulation by peptide docking studies below. All MD simulation output frame coordinates and binding energies are available on request.

### InteractiveROSETTA peptide docking to 50ns MD minimized states

As further orthogonal computational support for the coregulator peptide-docking studies, Frames #1, #2635, and #4719 identified from the shorter 50ns MD simulations described above were docked to peptides representing PGC1α, SHP, and DAX1 using InteractiveROSETTA^99^. The peptide sequences used for docking were identical to each peptide sequence used in wet lab *K*_d_ determinations by fluorescence polarization, except that no FAM-based fluorophores were present in the docked peptides. In each docking run, 1000 structures were generated, and the 10 refined models with lowest RMSD between output models were selected at end of each run for further analysis.

YASARA/FoldX^100^ was then applied to calculate binding energies and dissociation constants between LRH-1 and the coregulator peptides for the 10 refined output models for each docking run. The YASARA/FoldX calculated binding energies for all 10 output models was used to calculate an average energy and compared to the initial state of the MD simulation. The docked structure coordinates with the lowest calculated binding energy are shown, all docked position coordinates and binding energies are available on request.

### Luciferase reporter assays

The direct effect of bilirubin on the transcriptional activity of full length human LRH-1 was measured using dual luciferase-based reporter gene assay. Human embryonic kidney 293T (HEK293T) cells were maintained in DMEM with 10% FBS at 37°C under 5% CO_2_. Transient co-transfections were performed in bulk by plating 3 x 10^6^ cells per 10-cm plates with 9μg of total DNA (1:2.5:7.5 ratio of (1) Renilla luciferase under constitutive activation via a TK-based promoter: (2.5) firefly luciferase under the control of the *CYP7A1* promoter^36^: (7.5) Either full-length human LRH-1 controlled via CMV promoter, or an empty DNA plasmid) in a 1:3 DNA:PolyJet ratio. Transfected full-length human LRH-1 constructs encoded expression of 1) wild-type LRH-1, or 2) the A349F or 3) the A349W single point mutants of human LRH-1, generated from the wild-type construct using QuikChange mutagenesis (Agilent), all mutations were verified by sequencing. After 24 hours, all cells were replated in 384-well plates containing 50μM unconjugated bilirubin, vehicle control (100uM beta-cyclodextrin in DMSO) or no compound, at a density of 10,000 cells/well. Following an additional 20-hour incubation period, both firefly and Renilla luciferase activity were measured using Biotek luminescence plate reader. The fold-change in the activity of LRH-1 for bilirubin compared to vehicle was calculated using the equation FC = (Cyp7_bilirubin_ – Empty_bilirubin_) / (Cyp7veh – Empty_veh_), where Cyp7_bilirubin_ is the normalized luminescence (fLuc/RLuc) for cells that have been transfected with full-length LRH-1 (WT or A349F or A349W) in the presence of compound, Empty _bilirubin_ is the normalized luminescence (fLuc/RLuc) for cells transfected with the empty vector plasmid control DNA in the presence of bilirubin, Cyp7_veh_ is the normalized luminescence fLuc/RLuc for cells transfected with full length LRH-1 in the vehicle treated cells and Empty_veh_ is the normalized luminescence fLuc/RLuc for cells transfected with empty DNA plasmid for vehicle treated cells. All raw luciferase data are available on request.

### Construction of *SLCO1B1* stable HepG2 cells

Polyclonal HepG2 cells stably expressing the human full length *SLCO1B1* cDNA (NCBI accession: NM_006446.5) were generated by transfecting HepG2 cells (ATCC HB-8065) using Lipofectamine 3000 in a piggyback transposase-based pPB plasmid along with a plasmid expressing piggyback transposase and the pPB plasmids encoding either human *SLCO1B1* or empty vector. After 48 hours the cells were selected with 750ug/ml hygromycin reflecting a hygromycin kill-curve executed on these HepG2 cells mock transfected with Lipofectamine 3000. Stable transformants were selected for 3 weeks and pooled to create a polyclonal, stable cell line used in all HepG2 cell-based studies. This polyclonal cell line is freely available upon request.

### RNA sequencing of transcriptomes

Polyclonal, stable human HepG2 cells expressing *SLCO1B1* were seeded at 0.5X10^6^ cells in 3ml L-Glucose DMEM media supplemented with 1mg/mL glutathione, 10% FBS, 1% Pen/Strep and 250ug/mL hygromycin in a 6-well plate overnight at 37°C, then treated with 50uM bilirubin (16.4ul of a 9.18mM stock in 100uM beta-cyclodextrin in DMSO) in 3ml media or an equal volume of vehicle control (16.4ul of 100uM Beta-cyclodextrin in DMSO). Cells were incubated for 24hrs at 37°C, 5% CO2 in a cell culture incubator, and total RNA prepared using Quick-RNA MiniPrep (Zymo Research) according to manufacturer recommendations. DNase-treated total RNA was then used to generate PolyA-selected RNA-seq libraries, which passed quality control (QC) and were sequenced on a NovaSeq 6000 (Illumina), using 150bp paired-end sequencing with actual number of reads of at least 41 million reads per sample. QC of raw reads were tested using FastQC (0.11.9), low quality reads were removed and Illumina adapter sequences trimmed using bbDuk (https://jgi.doe.gov/data-and-tools/bbtools/bb-tools-user-guide/bbduk-guide/).

Trimmed sequences were then mapped to hg38 using HiSat2^101^. Poorly mapped and unaligned sequences were removed using Hisat2^101^ and SamTools^102^. Mapped reads were counted using feature counts^103^, raw read counts were used to assess differential expression using DESEQ2 with Apeglm LFCShrinkage in the R Bioconductor package. Differentially expressed gene sets were subjected to Gene Set Enrichment Analysis (GSEA) using the fgsea package in R bioconductor using the Molecular signatures database (version 7.4) as the reference. All original FASTQ files are available upon request.

### Reverse-transcriptase semi-quantitative polymerase chain reaction (RT-qPCR)

First strand cDNA was synthesized from 2ug total RNA using random-primed reverse transcription (Verso cDNA Synthesis, Thermo Scientific) using a thermal cycler according to manufacturer’s instructions. First strand cDNA synthesis was done at 42°C for 30min followed by 2 minutes inactivation at 95°C. Quadruple biological replicates with 2 technical replicates were used for each RT-qPCR condition, no data points were excluded from the analysis. *GAPDH* was the reference gene for normalization, and all primers were validated at serially diluted cDNA concentrations to produce a single PCR product in the dissociation curve. Primer sequences (all human genes) were: Cyclophilin (*PPIA*) Forward: 5’-GCGACTTTCTGGAGTTTATTTCA -3’; Cyclophilin (*PPIA*) Reverse: 5’-TTTCATTGCTTCTGGGTTCC -3’ *GAPDH* Forward: 5’-CAAGGTCATCCATGACAACTTTG-3’; *GAPDH* Reverse: 5’-GGCCATCCACAGTCTTCTGG-3’; *SLCO1B1* Forward: 5’-CGTAGAGCAACAGTATGGTCAGC-3’; *SLCO1B1* Reverse: 5’-TTGGCAATTCCAACGGTGTTCAG-3’; *NR0B2* Forward: 5’-GCTTAGCCCCAAGGAATATGC-3’; *NR0B2* Reverse: 5’-TTGGAGGCCTGGCACATC-3’; *CYP7A1* Forward: 5’-GCGACTTTCTGGAGTTTATTTCA-3’ *CYP7A1* Reverse: 5’-TTTCATTGCTTCTGGGTTCC-3’; *CYP11A1* Forward: 5’-GGGTCGCCTATCACCAGTATT-3’; *CYP11A1* Reverse: 5’-GCTGCCGACTTCTTCAACAG-3’; *CYP17A1* Forward: 5’-AGGACTTCTCTGGGCGGCCT-3’ *CYP17A1* Reverse: 5’-GTGTGCGCCAGAGTCAGCGA-3’ *NR5A2* Forward: 5’-CAGAGAAAGCGTTGTCCTTACTG-3’; *NR5A2* Reverse: 5’-TTATTCCTTCCTCCACGCATT-3’. CFX96 Real Time qPCR instrument (BioRad, Hercules, CA) was used for real time data acquisition using the following cycling parameters: pre-denaturation 95**°**C for 5 minutes, denaturation 95**°**C for 10 sec, annealing and extension 60**°**C for 30sec for a total of 40 cycles. A melt curve step at 65**°**C for 5 sec and 56**°**C for 50 sec in every run further validated the specificity of primers to produce only one PCR product. The ΔΔCt method using a reference gene (Cyclophilin *PPIA* or *GAPDH*) Ct subtracted from the Ct of the target gene of interest normalized transcript levels for each cDNA sample. The data were analyzed in GraphPad Prism by one-way ANOVA with Dunnett’s correction for multiple comparisons from all quadruplicate biological and technical duplicates. All raw qPCR data are available on request.

### Transcriptome enrichment in ChIP gene sets

Published chromatin immunoprecipitation (ChIP) gene sets, all collected in human HepG2 cells, were obtained from published sources, identifying the direct chromatin target genes for LRH-1^61^, AHR^64^, PPARα^65^, PPARγ^63^ and SP1^63^ as provided in Supplemental Spreadsheet 2. Quantifiable genes in each transcriptome were defined as all transcripts with any *p* value assigned by DESEQ2, percentage of bilirubin regulated genes in the quantifiable transcriptome was used as the expected discovery rate for continency analyses (Supplemental Spreadsheet 3). Quantifiable ChIP genes for LRH-1, AHR, PPARα, PPARγ and SP1 were defined as any gene in the ChIP gene set present as a quantifiable gene in the transcriptome, and are provided in Supplemental Spreadsheet 2. Enrichment of bilirubin-regulated genes in all quantifiable genes *vs.* the quantifiable ChIP-seq gene set were determined by Fisher’s exact tests using GraphPad Prism, and are provided in Supplemental Spreadsheet 3. Enrichment of bilirubin-regulated genes in the LRH-1 ChIP-seq gene set were compared to enrichment in ChIP gene sets from AHR, PPARα, PPARγ or SP1 individually to LRH-1, not simultaneously. All contingency analyses were done in GraphPad Prism, the quantifiable gene sets are provided as supplemental spreadsheets.

### Small interfering RNA knockdown of LRH-1 (NR5A2) transcriptome

HepG2 cells stably expressing *SLCO1B1* were seeded (0.5X10^6^ cells) in 3ml L-Glucose DMEM media supplemented with 1mg/mL glutathione, 10% FBS, 1% Pen/Strep and 250ug/mL hygromycin (to maintain the *SLCO1B1* expression cassette) in a 6-well plate overnight at 37°C, 5% CO_2_. The following morning, cells were trypsinized and reverse transfected into a new 6 well plate with 20nM siLRH-1 per well (ON-TARGETplus Human *NR5A2* (2494) siRNA-SMARTpool, 10nmol, Dharmacon L-003430-00-0010), or 20nM siPPARa (ON-TARGETplus Human *PPARA*), 20nM siPPARg (ON-TARGETplus Human *PPARG*), 20nM siSP1 (ON-TARGETplus Human *SP1*) or 10nM siCON per well (non-targeting siRNA, On-TARGETplus Non-targeting Pool, Dharmacon D-001810-10-05) with Dharmafect transfection reagent (Horizon). After 72 hours siRNA, wells were treated with either 50uM unconjugated bilirubin or vehicle alone (100uM Beta-cyclodextrin in DMSO). Cells then were incubated for another 24 hours (96 hours total siRNA, 24 hours total bilirubin or vehicle), then total RNA extracted (Quick-RNA MiniPrep, Zymo Research), DNase treated RNA samples used for polyA-selected RNA library preparation and sequencing, as detailed above. All original FASTQ files are available upon request.

### RJW100-competitive transcriptome

RJW100^53^ is a mixture of both *RR* and *SS* enantiomers and was synthesized by the Vanderbilt Institute for Chemical Biology according to the published synthesis^53^ but is also commercially available from several sources. Both enantiomers in the mixture have been shown to bind the canonical ligand-binding site of LRH-1 and competitively displace other ligands from this site^53^. HepG2 cells (0.5X10^6^ cells) stably expressing *SLCO1B1* bilirubin transporter were seeded in 6 well plate overnight, were then co-treated at the same time with Vehicle + 100uM RJW100; 50uM bilirubin + 100uM RJW100; or the appropriate vehicle alone controls, the vehicle for RJW100 was DMSO, the vehicle for bilirubin is 100uM cyclodextrin in DMSO. Total RNA was prepped after 24 hours compound treatment using Quick-RNA MiniPrep (Zymo Research) according to manufacturer recommendations and used for polyA-selected RNA library preparation and sequencing as detailed above. All original FASTQ files are available upon request.

## List of Supporting Information

**Supplemental Figures S1-S14:** Supporting data for main text figures.

**Supplemental Tables 1-2**: Cluster analysis of 25,000 MD simulated states of the bilirubin/LRH-1 complex

**Supplemental Spreadsheet 1:** Bilirubin *vs*. Vehicle transcriptome results

**Supplemental Spreadsheet 2:** Quantifiable ChIP gene sets

**Supplemental Spreadsheet 3:** Enrichment of bilirubin-regulated genes in quantifiable ChIP gene sets

**Supplemental Spreadsheet 4:** Bilirubin+siCON *vs*. Vehicle+siCON transcriptome results

**Supplemental Spreadsheet 5:** Bilirubin+siLRH-1 *vs*. Vehicle+siLRH-1 transcriptome results

**Supplemental Spreadsheet 6:** Vehcile+RJW100 *vs*. Vehicle+DMSO transcriptome results

**Supplemental Spreadsheet 7:** Bilirubin+RJW100 *vs*. Vehicle+RJW100 transcriptome results

**Supplemental Spreadsheet 8:** Energies of 2000 Rosetta poses, bilirubin docked to LRH-1

**Supplemental Spreadsheet 9:** MD simulation (0.5us) RMSF and RMSD results

**Supplemental Spreadsheet 10:** MD simulation (50ns) RMSF results

**Supplemental Spreadsheet 11:** MD simulations (50ns) RMSD results

**Supplemental Spreadsheet 12:** Bilirubin *vs*. Vehicle Gene Set Enrichment Analysis (GSEA) results

## Acknowledgments

This work was supported by the National Institutes of Health GM138873 to RDB. ZH was supported by the Vanderbilt Genomic Medicine Training Program T32HG008341. PC was supported by Vanderbilt Molecular Endocrinology Training Program DK007563. RWJ100 was provided by the Molecular Design and Synthesis Center (MDSC) and the VICB Discovery Collection was distributed and screened by the Vanderbilt High-throughput Screening (HTS) Core Facility. The Center for Innovative Technology at Vanderbilt University, HTS and MDSC receive support from the Vanderbilt Institute of Chemical Biology and the Vanderbilt Ingram Cancer Center P30 CA68485.

---

**Supplemental Figure S1:**
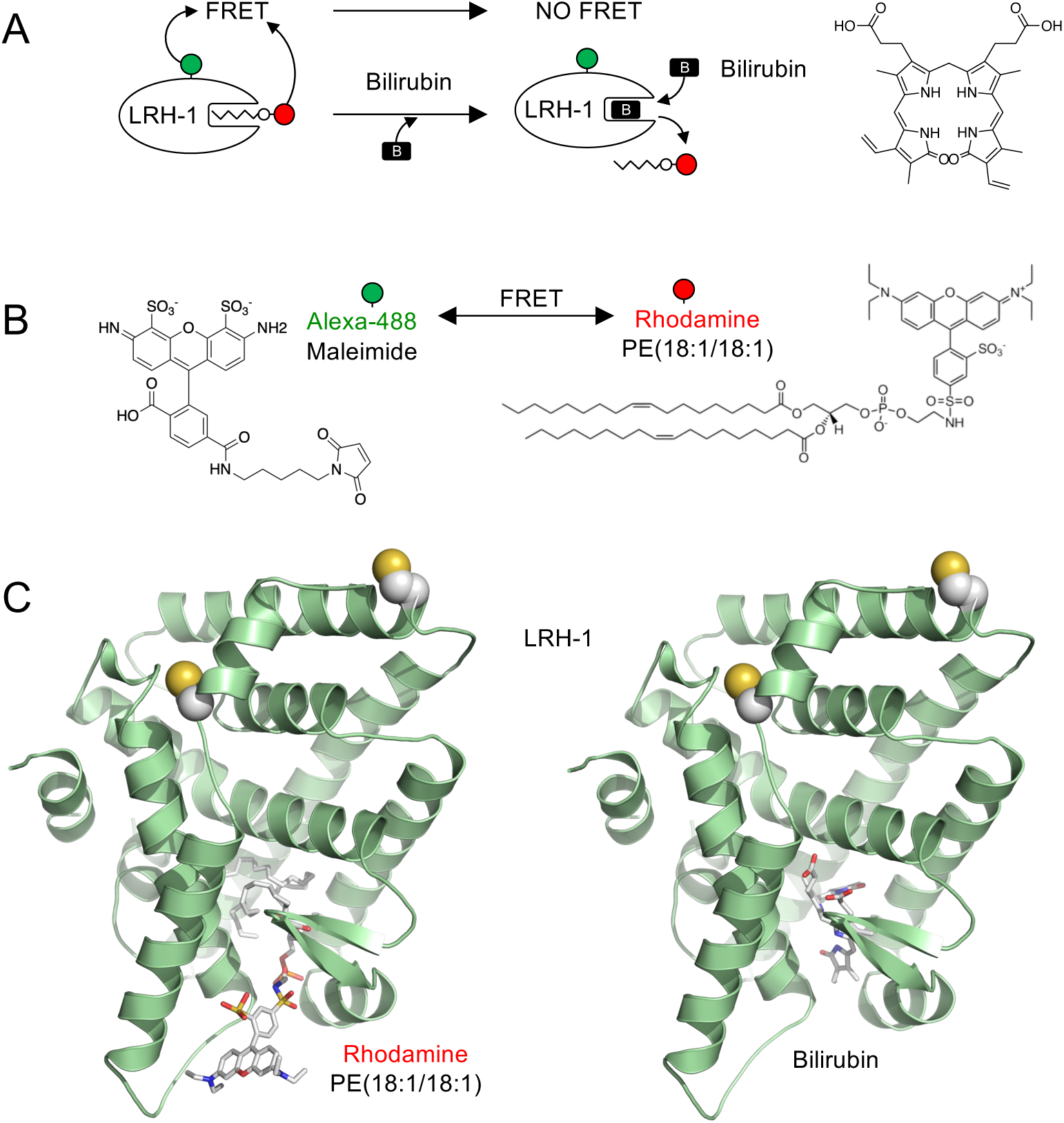
LRH-1 displacement binding assay design (adapted from Malabanan, et al. ACS Chemical Biology, 2023). **A**. Schematic of the assay design, showing bilirubin biding to the canonical, orthosteric ligand binding site in LRH-1 should displace the phospholipid-based Rhodamine Phosphatidylethanolamine (Rh-PE) probe. **B**. Chemical structures of label for LRH-1 (Alexa-488 Maleimide), which chemically reacts with free Cys residues, and the Rh-PE probe (Avanti Polar Lipids). **C**. PyRx rigid-body computational docking of Rh-PE and bilirubin to the crystal structure of human LRH-1 ligand binding domain (PDB:60QX) with two maleimide-reactive Cys side chains indicated as spheres, suggesting both Rh-PE and bilirubin are capable of binding to the same site in LRH-1. These schematics and docking experiments clarify the design of the direct binding assay used to obtain the apparent *Kd* of bilirubin for LRH-1 reported in Figure 1.

**Supplemental Figure S2:**
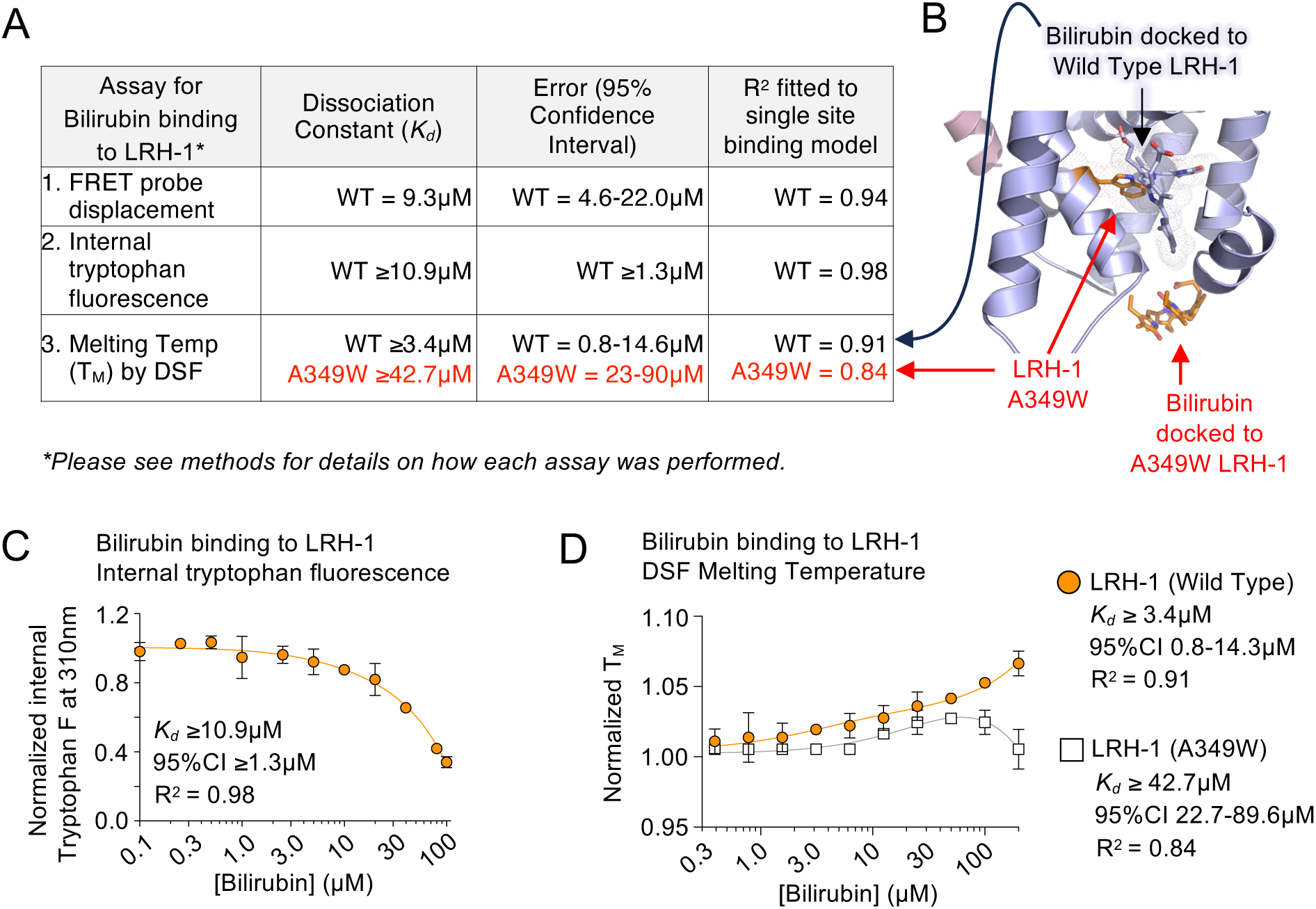
Three independent assays confirm bilirubin Kd for LRH-1 of approximately 10µM. **A.** Compiled table of 3 independent assays measuring the dissociation constant of bilirubin for the purified ligand binding domain of LRH-1, error represents 95% CI as indicated, red indicates data collected for the pocket mutant of LRH-1 (A349W), previously shown to bind ligand poorly, if at all. **B**. PyRx models of bilirubin docked to either WT LRH-1 (purple) or A349W mutant LRH-1 (orange, labels in red), illustrating the predicted steric clash between mutant A349W residue and bilirubin. Bilirubin is not predicted to bind in the canonical ligand binding pocket of A349W mutant LRH-1 (PDB:6OQX). **C**. Curve fit of internal tryptophan fluorescence data of bilirubin binding to purified wild-type LRH-1 ligand binding domain, upper limit of 95% CI could not be defined, error represents 95% confidence interval. **D**. Curve fits of normalized LRH-1 melting temperature (TM) changes induced by increasing concentrations of bilirubin, as determined by Differential Scanning Fluorimetry (DSF). Note that both wild type (WT) and the pocket mutant (A349W) were used to determine the binding of bilirubin to LRH-1, and that the pocket mutant had worse interaction qualities with bilirubin when compared to wild-type LRH-1. These data suggest that in multiple, independent assays, bilirubin binds directly to the LRH-1 ligand binding domain with Kd ∼ 10µM, within the 95% confidence intervals of all three assays.

**Supplemental Figure S3:**
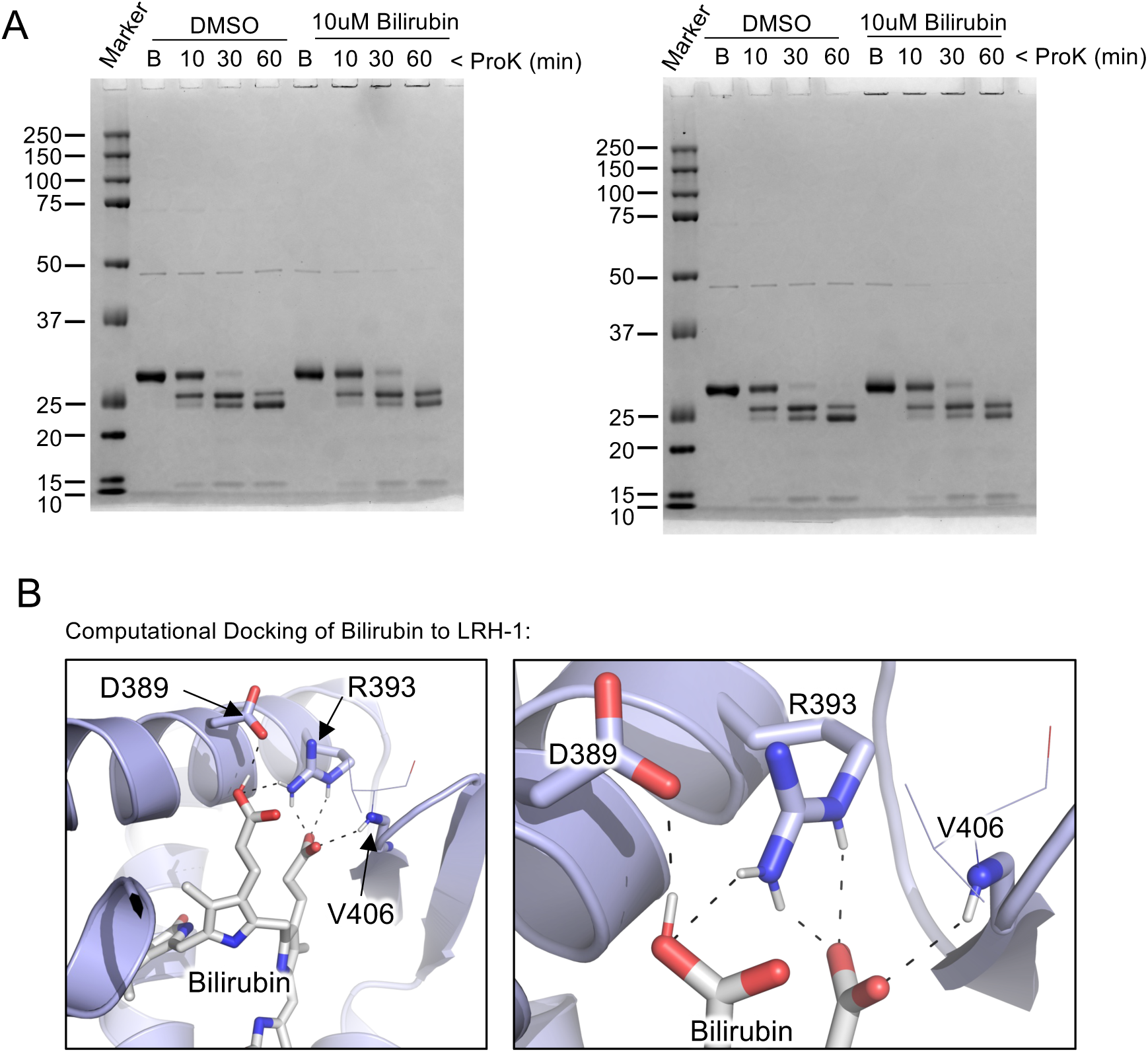
Bilirubin stabilizes LRH-1 structure. **A**. Full gel images of Coomassie-stained SDS-PAGE of proteinase-K LRH-1 ligand binding domain protection induced by overnight incubation with bilirubin at the indicated times of ProK digestion at 25C. “B” represents Buffer only no Pro-K added sample incubated at 25C for 60 minutes as negative control in the presence of indicated DMSO or bilirubin. ANOVA analysis of the top band representing full-length, undigested LRH-1 LBD showed significant ProK protection of LRH-1 induced by bilirubin across all the time points (*p*=0.0458), statistical analyses done in GraphPad Prism. **B**. PyRx rigid-body docked positions of bilirubin to the LRH-1 LBD, showing potential contacts with D389, R393 and V406, suggesting one of many potential binding positions that are plausible *in silico*. These data suggest bilirubin slows Proteinase-K digestion of the purified LRH-1 ligand binding domain, *in vitro*.

**Supplemental Figure S4:**
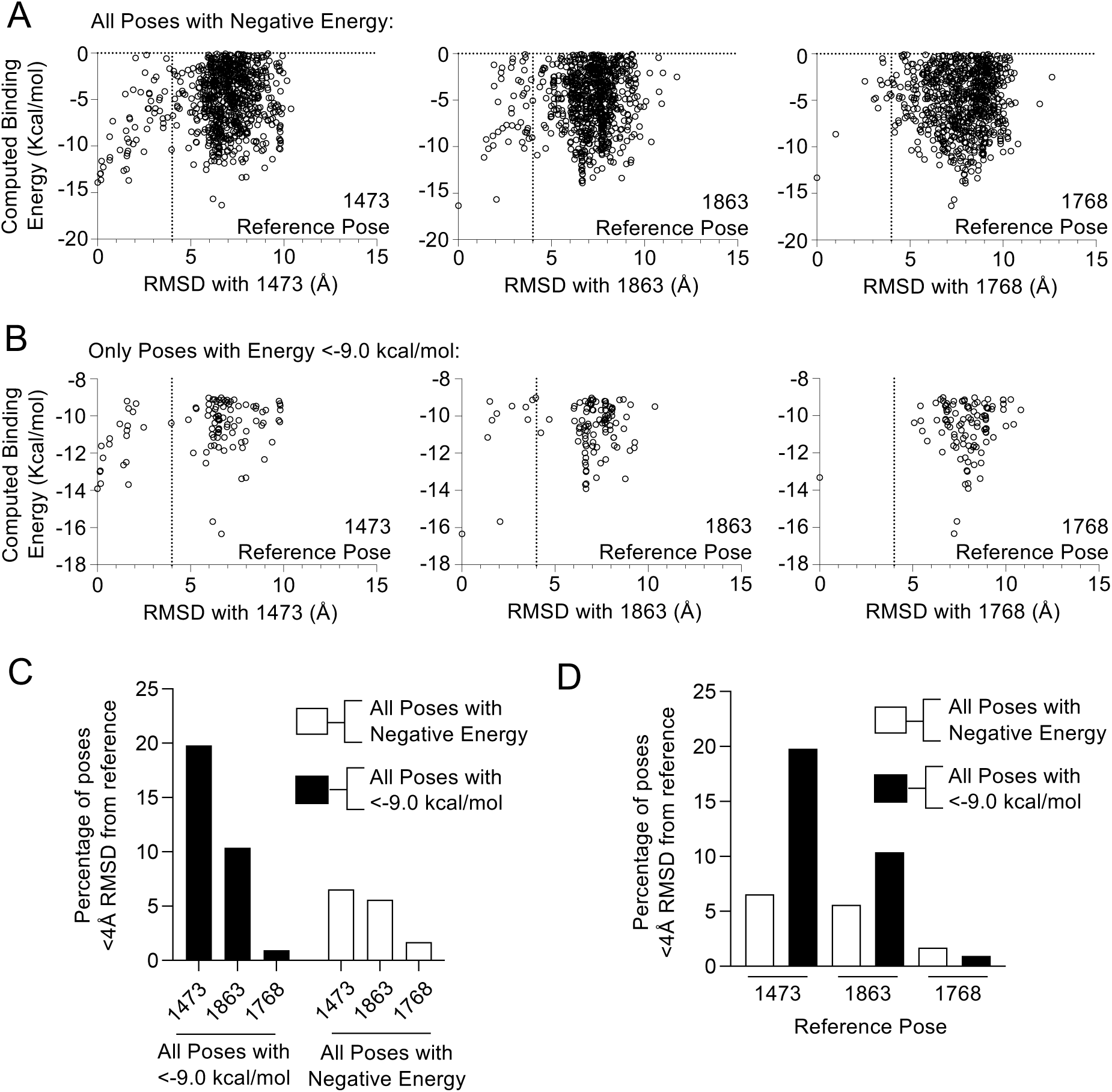
Rosetta docking of bilirubin to LRH-1 produced 3 groups of similar poses, pose 1473 is the lowest energy pose in the most frequently populated group. Bilirubin was docked to LRH-1 using Rosetta to produce 2000 unique poses of bilirubin bound to LRH-1. The ten poses with lowest energy from these 2000 were used as reference poses to calculate the root mean square deviation (RMSD) with all 2000 poses. This resulted in 3 groups of similar poses, the most frequently populated of these three groups contained pose 1473, which was the lowest energy pose with the group. **A**. All poses with negative energy were plotted as a function of bilirubin binding energy (kcal/mol) and RMSD (Å) from indicated reference pose (1483, 1863 or 1768). Line indicates 4Å RMSD from indicated reference poses, used for analysis in C. & D. **B**. All poses with binding energy below -9kcal/mol were plotted as a function of bilirubin binding energy (kcal/mol) and RMSD (Å) from indicated reference poses (1483, 1863 or 1768). Established wet-lab LRH-1 high affinity reference compounds 5N and 6N both bound with calculated binding energies of -9.0 kcal/mol, thus establishing this value as a stringent threshold for these computational experiments. Line indicates 4Å RMSD from indicated reference poses. **C.** Frequency of poses (percentage or number of poses per 100 poses) that occur with less than 4Å RMSD from the indicated reference poses. Note the 1473 group has the highest frequency of low-RMSD poses, regardless of which group was analyzed. **D**. Same analysis as in C. but grouped by reference pose. These data suggest pose 1473 is the lowest-energy and most frequently populated pose of bilirubin bound to LRH-1 as determined by Rosetta. Pose 1473 was selected for further MD simulation analysis.

**Supplemental Figure S5:**
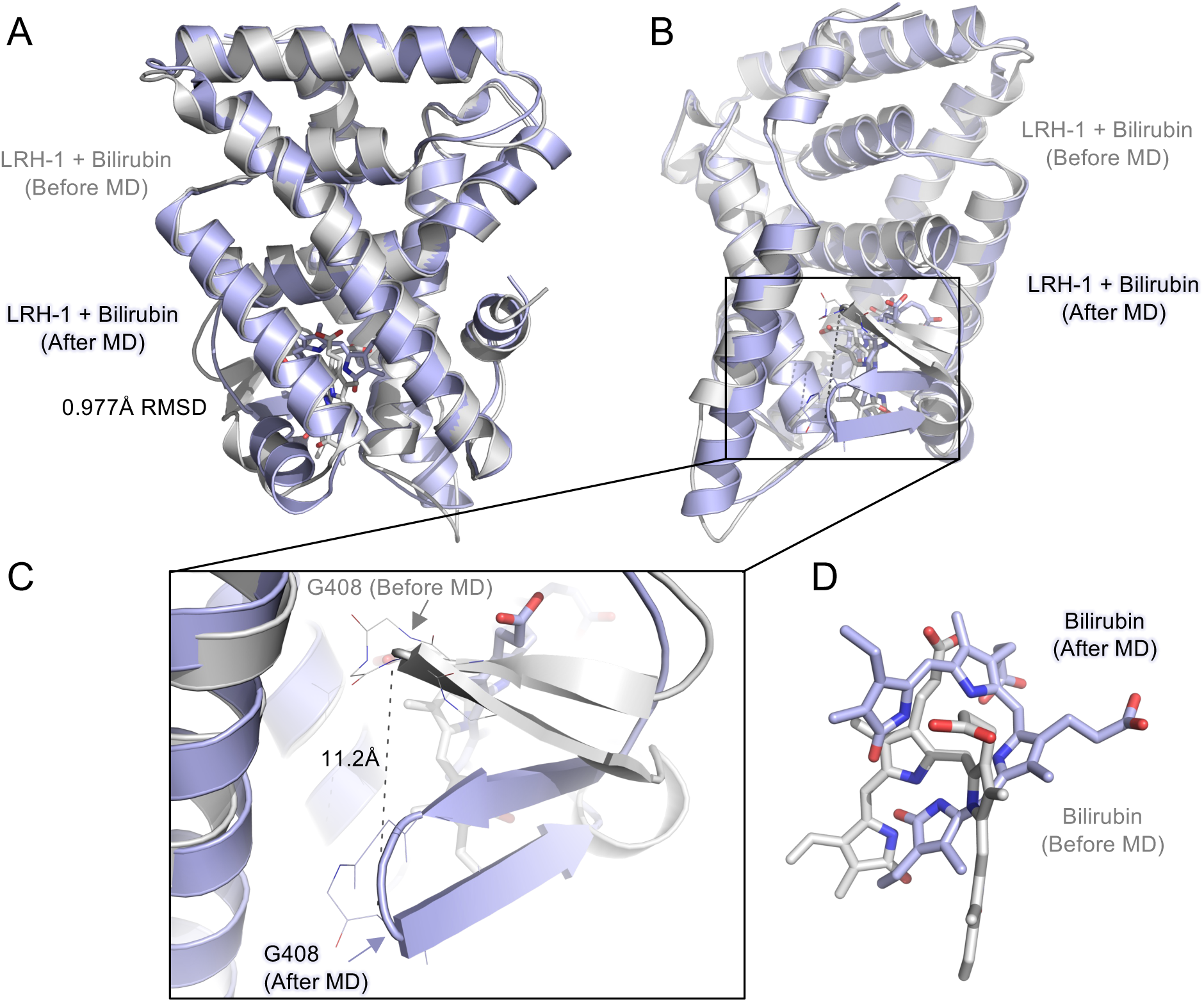
0.5µs MD simulation suggests bilirubin stabilizes an alternative conformation of LRH-1. Rosetta docking of bilirubin to LRH-1 produced 2000 poses, of which pose number 1473 was the most frequently populated. Pose 1473 was then used as time zero (T0) in 0.5µs MD simulations. Results from MD were analyzed according to Table 1. **A**. Cartoon representations of LRH-1 bound to bilirubin in the initial state (“Before MD”, pose 1473, cartoon grey) superposed with the most frequently populated state from the MD simulation (“After MD”, cartoon blue), which is frame 15783 from 1473-1 cluster representing the centroid structure from from the most populated cluster of states from the 0.5µs MD simulation (see Table 1 for clustering results). **B**. Same is in A, but rotated 180°, box drawn around the beta sheet of the AF-B regulatory region of LRH-1. **C**. Detail of boxed region, showing 11.2Å shift in residue G408 after MD simulation. Once attaining this position, the AF-B beta-sheet region remained stable in this position for the final 460ns of the MD simulation. **D**. Stick representation of bilirubin molecules with LRH-1 protein cartoons removed at T0 (grey sticks) compared to the most frequency populated state from the MD simulation (blue sticks, frame 15783 from 1473-1 cluster, see Table 1). These computational representations illustrate one way bilirubin is predicted to alter LRH-1 conformation, and further highlight that bilirubin binding to the canonical ligand binding site is stable *in silico*.

**Supplemental Figure S6:**
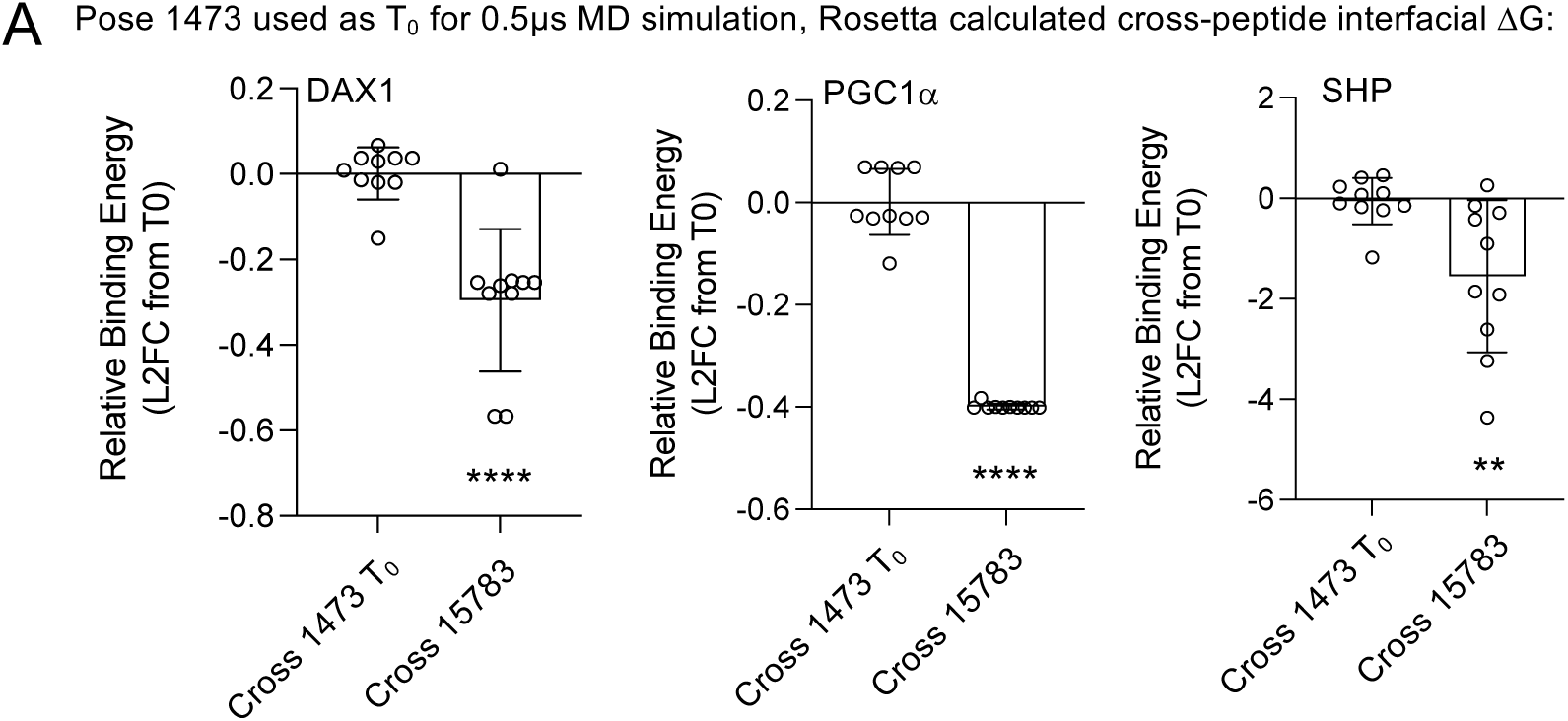
Bilirubin regulates LRH-1 interaction with transcriptional co-regulator peptides *in silico*. We used Rosetta to dock bilirubin to LRH-1, producing 2000 poses, of which pose 1473 represented the lowest energy pose in the most frequently populated group of poses. We used pose 1473 as the initial state (T_0_) in 0.5µs MD simulation, and applied clustering to determine the most frequently populated of 25,000 MD states. We found frame **15783** was the centroid structure from the most populated cluster from the MD simulation (cluster #1473-1, see Table 1). Rosetta was then used to compute the interfacial binding energy between 3 separate transcriptional coregulator peptides and the LRH-1/bilirubin complex, both in the initial state before MD simulation, (1473 T0) and after MD simulation (frame 15783). **A**. Log2 fold change of the computed energy of indicated coregulator peptides docked to either the T0 initial state before MD (pose 1473, before MD), or the centroid frame of the most frequently populated MD state (frame 15783, after MD), expressed relative to the initial state. Structures used for peptide docking here are identical to those used in Fig 6G, but here energy of the peptide docked to the LRH-1/bilirubin complex was measured as the cross-peptide interfacial *ΔG* (dG_cross), or energy of the interface calculated with cross-interface terms. Fig 6G *ΔG* is the difference between the peptide bound vs. unbound states (dG_separated), calculated by separating and repacking LRH-1 and coregulator peptide chains. Each point is one of 10 Rosetta peptide docking runs, error represents standard deviation, all t-tests were Bonferroni-corrected for multiple comparisons (\*\**padj*<0.01, \*\*\*\**padj*< 0.0001) as docking runs were computed simultaneously. These data suggest bilirubin stabilizes LRH-1 into a conformation that regulates LRH-1 interaction with coregulator peptides, regardless of how the peptide-LRH-1 interface is analyzed, also supported by wet lab data presented in Figure 1.

**Supplemental Figure S7:**
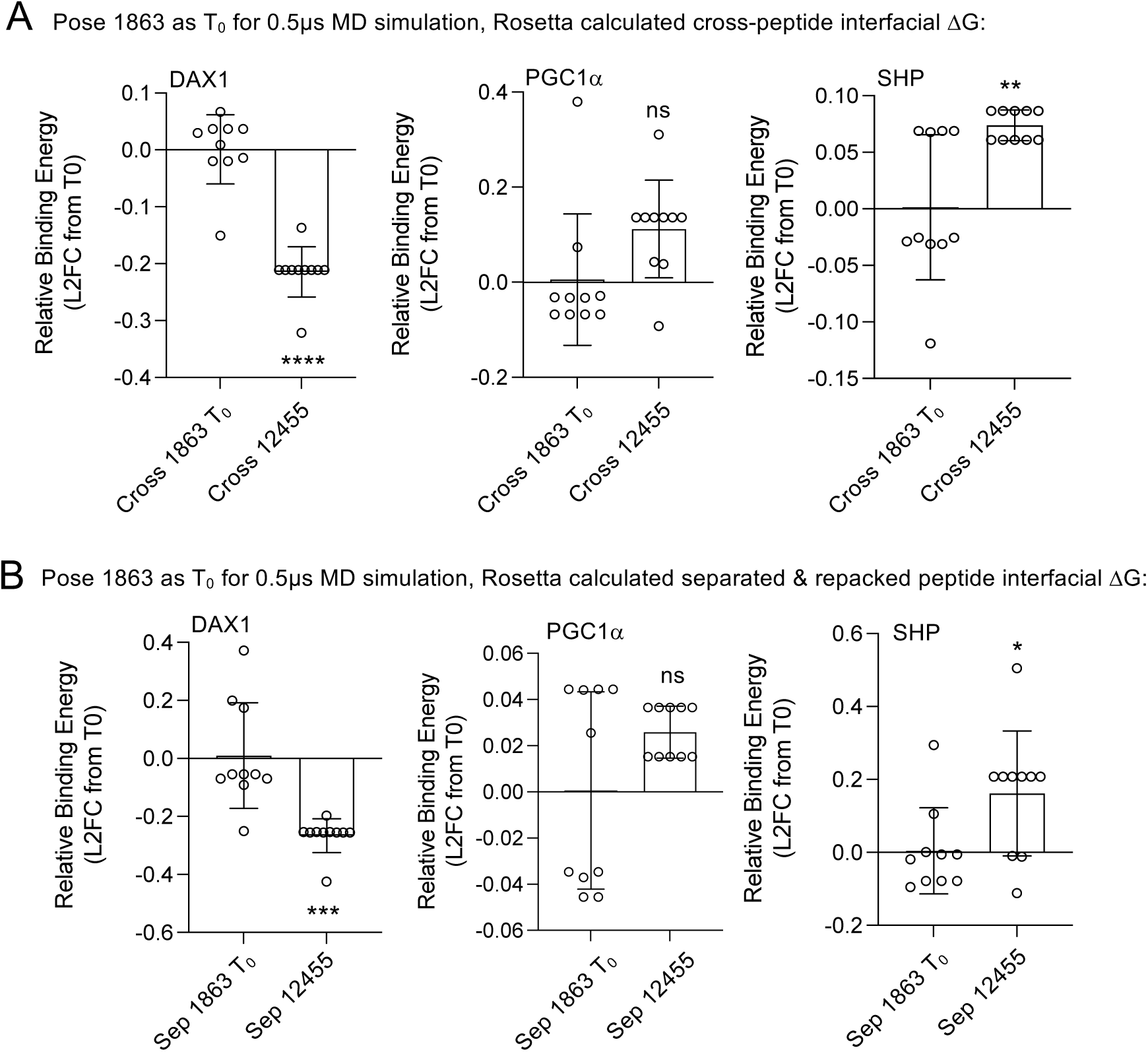
Bilirubin regulates LRH-1 interaction with transcriptional co-regulator peptides in silico. Rosetta docking of bilirubin to LRH-1 produced 2000 poses of which pose 1473 was the most frequently populated pose, while a second pose (1863, analyzed here) was the second most frequently populated pose, and was used as the initial state (T0) in a second independent 0.5µs MD simulation. Pose 1863 was used as the initial state (T0) in Rosetta-based computational docking to coregulator peptides, along with the centroid from the most populated state from the 0.5µs MD simulation (Frame **12455** from cluster 1863-1, see Table 2). Frame 12455 represents the output of the MD simulation in the presence of bilirubin, when the simulation was initiated with pose 1863. **A**. Log2 fold change of the dG_cross binding energy (*ΔG*) of indicated coregulator peptides docked to frame 12455 (the centroid of the most frequently populated MD cluster using pose 1863 as T0). **B**. Log2 fold change of the dG_separated binding energy (*ΔG*) of indicated coregulator peptides docked to pose 1863 (before MD) or frame 12455 (after MD). These data suggest that MD simulation using pose 1863 as the initial state generated LRH-1 conformations that less dramatically regulate LRH-1 interactions with coregulator peptides. In total, the MD simulations with bilirubin produced conformations which regulated LRH-1 interaction with coregulator peptides in 12/15 *in silico* tests. Each data point shown represents one of 10 different peptide docking runs, error represents standard deviation between runs, and all t-tests were Bonferroni-corrected for multiple comparisons (\*\**padj*<0.01, \*\*\*\**padj*< 0.001, \*\*\*\**padj*< 0.0001) as the docking runs were computed simultaneously. These data suggest bilirubin regulates LRH-1 interaction with coregulator peptides, also supported by the wet lab data presented in Figure 1.

**Supplemental Figure S8:**
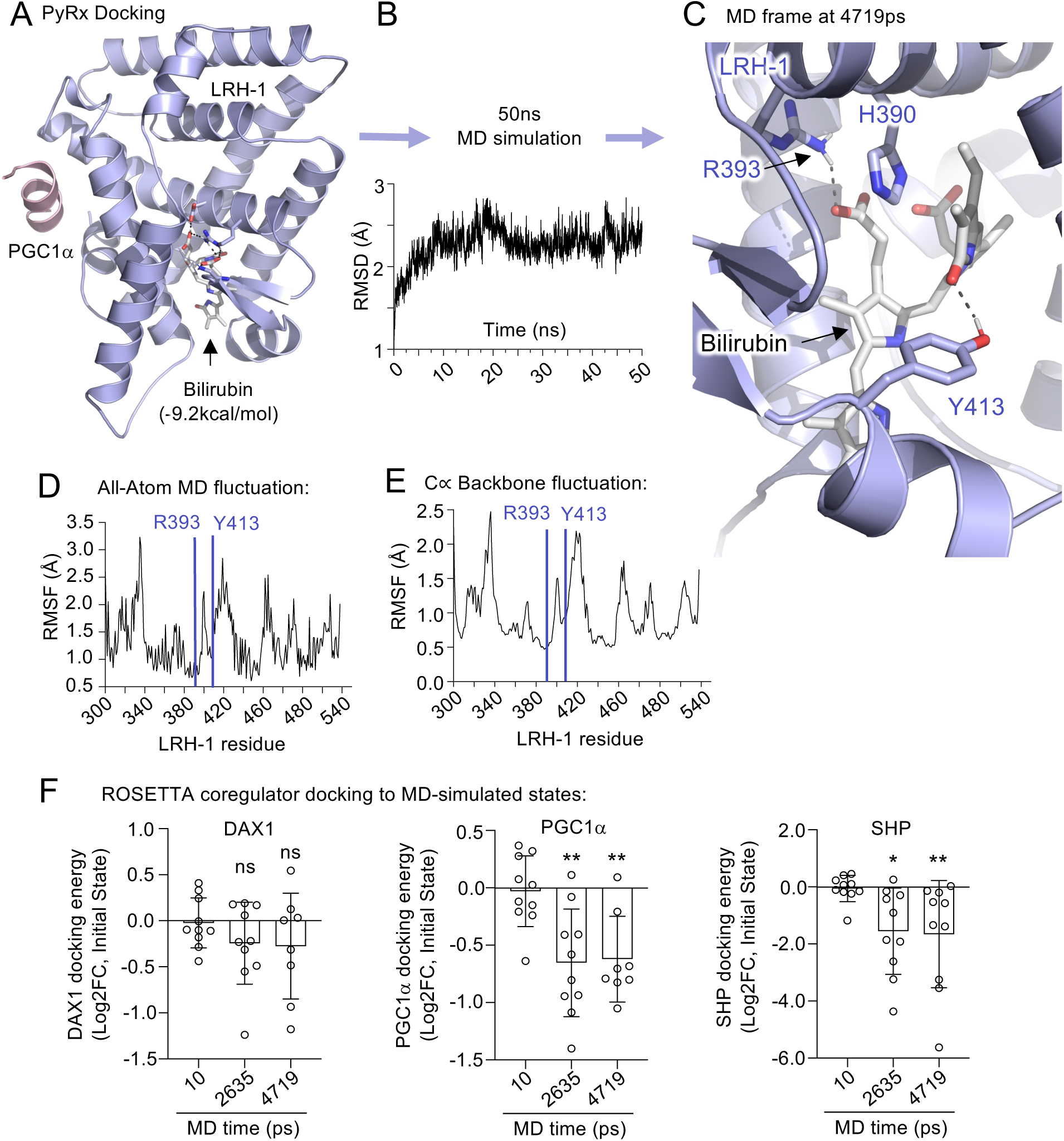
PyRx docking and 50ns MD simulations predict bilirubin binds stably to the canonical ligand binding site in LRH-1, and alters LRH-1 interaction with transcriptional coregulator peptides. **A**. Lowest energy PyRx rigid-body docked position from 9 output frames of bilirubin docked to the crystal structure of LRH-1 LBD (PDB:6OQX). This lowest energy frame was used as the initial state in **B**. 50ns AMBER molecular dynamics (MD) simulations, the calculated all-atom root mean square deviation (RMSD) is shown over MD time. Bilirubin remained stably bound to LRH-1 throughout the entire simulation. **C**. A closeup of the MD state after 4719ps, showing predicted interactions of bilirubin with R393 and Y413. **D**. All-atom root mean square fluctuation (RMSF) and **E**. C-alpha backbone RMSF of indicated LRH-1 amino acids over the 50ns MD simulation, R393 and Y413 indicated. **F**. Log2 fold change of the Rosetta-computed binding energy of indicated coregulator peptides docked to the lowest energy MD state, relative to the initial state in the MD simulation. Each point represents one of 10 independent Rosetta docking runs, error represents standard deviation between runs, all t-tests Bonferroni-corrected for multiple comparisons (\**p_adj_*<0.05, \*\**p_adj_*< 0.01). These data are consistent with bilirubin stably binding LRH-1 and regulating LRH-1 structure in a manner that alters LRH-1 interaction with transcriptional coregulator peptides, *in silico,* also supported by the wet lab data.

**Supplemental Figure S9:**
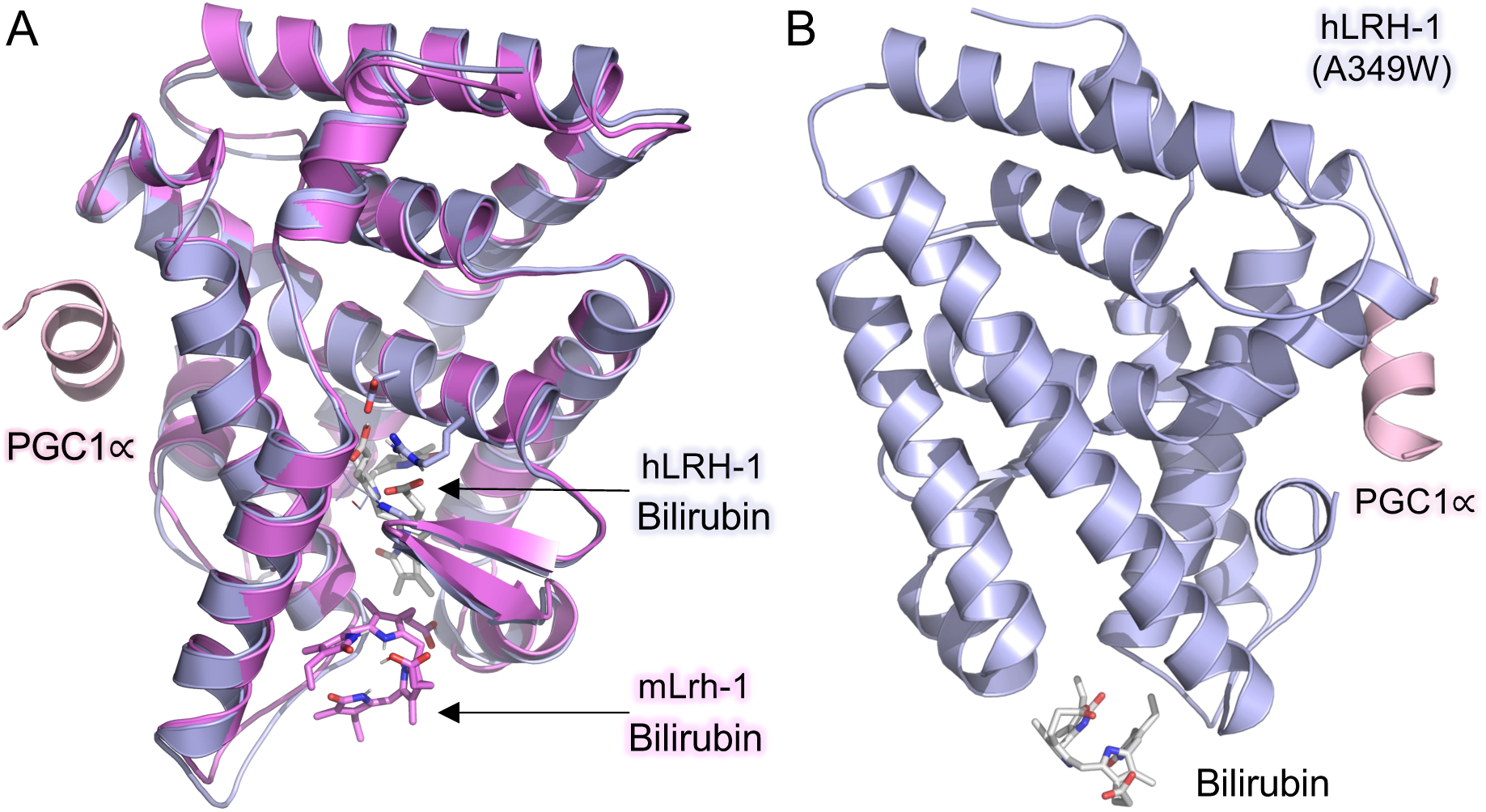
Bilirubin is not predicted to bind the mouse orthologue of LRH-1, or the A349W pocket mutant of human LRH-1. **A**. Superposed structures of bilirubin docked to either human LRH-1 (blue) or mouse Lrh-1 (magenta, PDB:1PK5) using PyRx rigid body docking. Note bilirubin cannot bind the canonical ligand binding site of mouse Lrh-1, PCG1∝ peptide shown in pink. **B**. PyRx docking result to a model of human LRHz-1 bearing the pocket mutant mutation (A349W), showing this mutant of human LRH-1 is not predicted to bind bilirubin in the canonical ligand binding site *in silico*. These data suggest that neither the mouse Lrh-1 orthologue nor the human A349W mutant are predicted to be sensitive to bilirubin.

**Supplemental Figure S10A-B:**
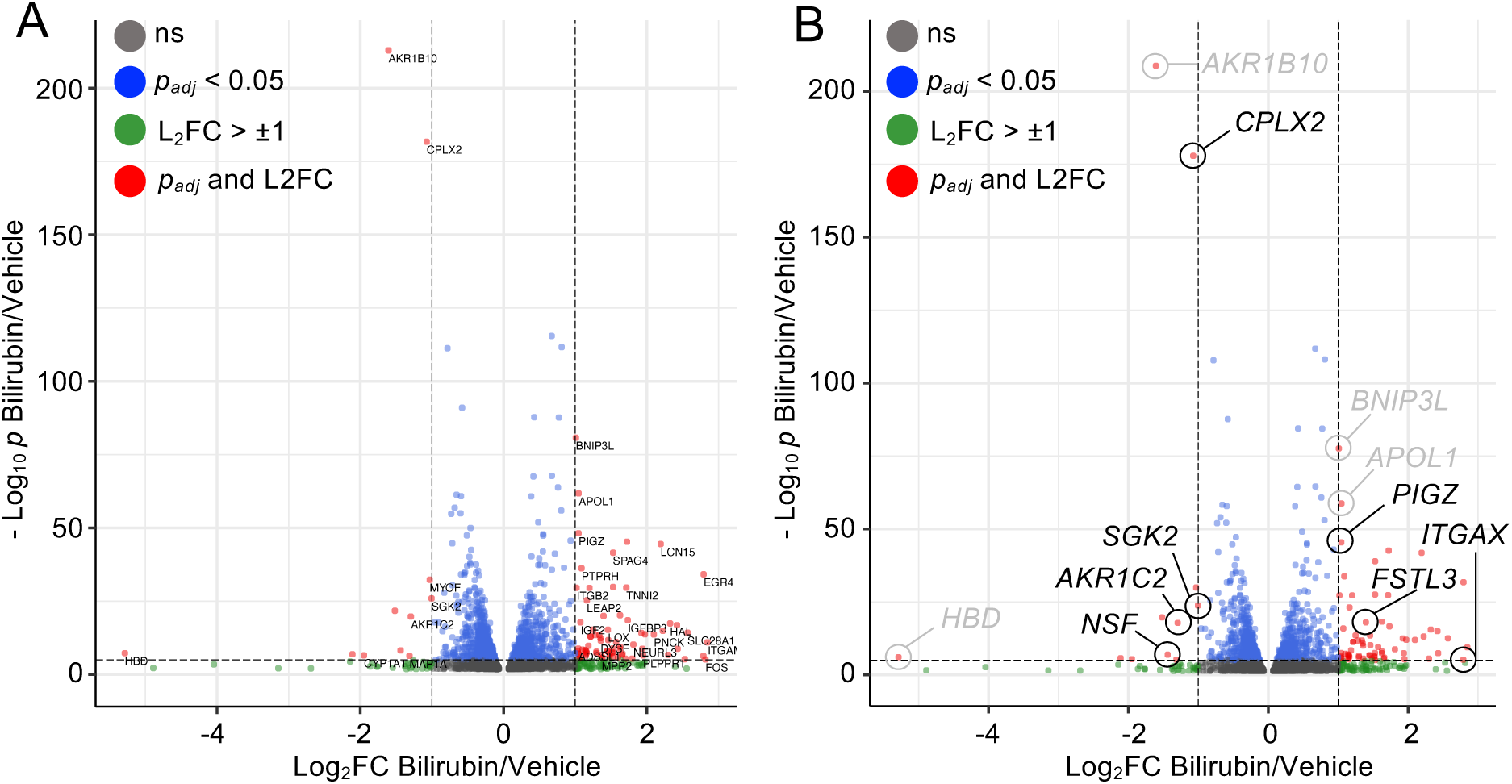
The bilirubin-regulated transcriptome in HepG2 cells is enriched in direct LRH-1 target genes. Both panels contain the same data with different labels for clarity. **B.** Full volcano plot of RNA-seq data from HepG2 cells treated with 50uM Bilirubin or vehicle control for 24 hours, presented to show the complete volcano plote, all regulated genes provided in supplemental spreadsheets. **C**. Same as in B., except selected direct LRH-1 ChIP-seq target genes labeled in black, transcripts labeled in gray are bilirubin regulated genes that were not identified in LRH-1 ChIP-seq studies. These data suggest that some, but not all, bilirubin regulated genes have been previously identified as direct LRH-1 ChIP-seq target genes.

**Supplemental Figure S11:**
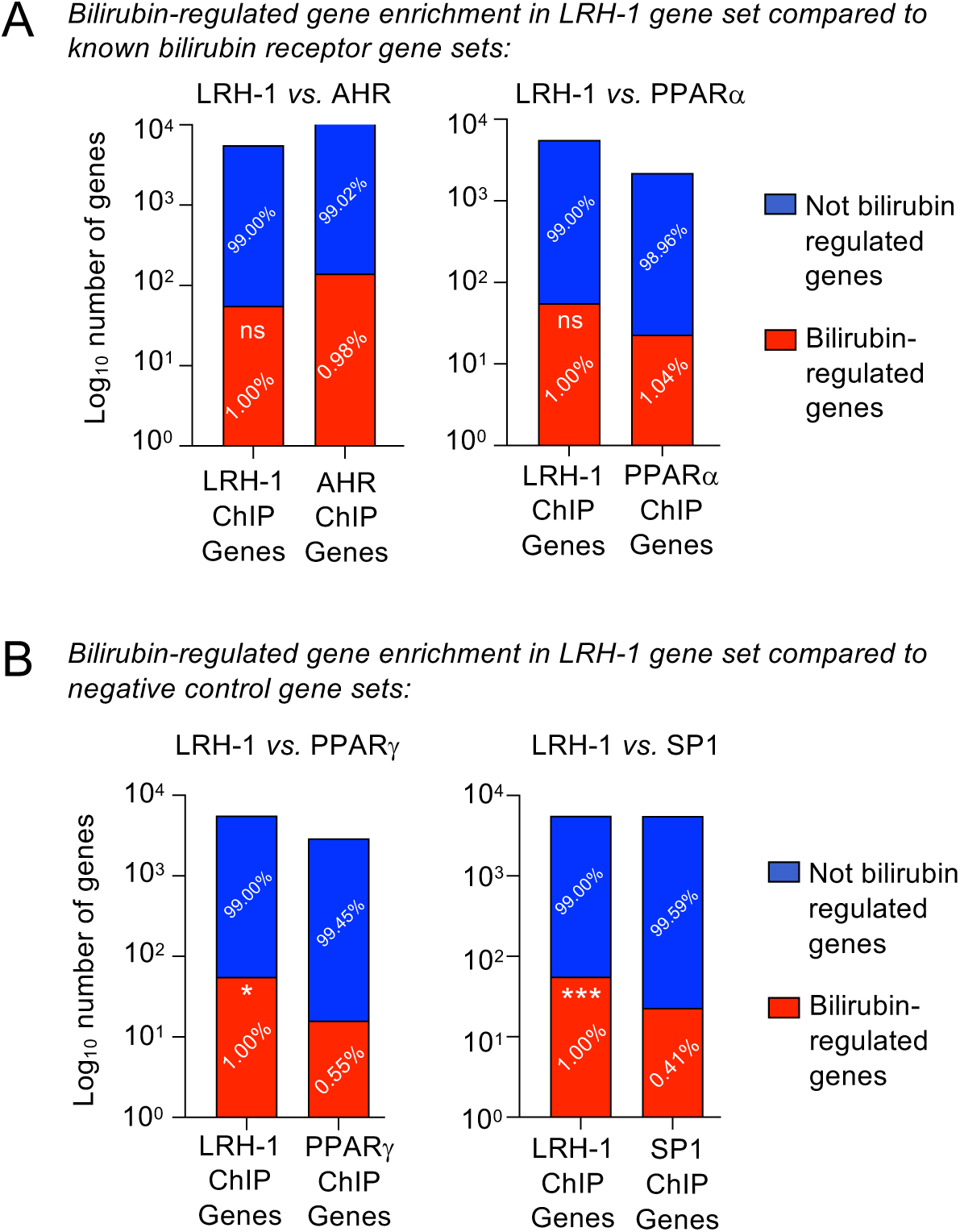
Direct LRH-1, AHR and PPARα target genes are similarly enriched in bilirubin-regulated genes. Contingency analyses plotting the Log10 number of genes regulated by bilirubin (red) and not regulated by bilirubin (blue) within each indicated ChIP gene set, with the percentages of each gene set also indicated. **A**. Number of bilirubin-regulated genes in the LRH-1 ChIP gene set from HepG2 cells, compared to the number of bilirubin-regulated genes in the AHR ChIP gene set also from HepG2 cells. Indicated percentages represent the percentage of bilirubin-regulated genes in each gene set (LRH-1 or AHR) and shows no significant difference (ns, not significant by Fisher’s exact test) in enrichment of bilirubin-regulated genes in the LRH-1 vs. AHR (left) or PPARα (right) ChIP gene sets. AHR and PPARα are known bilirubin receptors, these data suggest bilirubin regulates LRH-1 target genes just as much as AHR or PPARα target genes. All ChIP gene sets represent only those genes that were detectable (quantifiable) in the HepG2 transcriptomes (see methods for details). **B.** Same analysis as in A, but now comparing negative control ChIP-seq gene sets for transcription factors not suspected to bind bilirubin, using PPARγ (left) and SP1 (right) ChIP-seq gene sets from ENCODE in HepG2 cells, showing statistically significant enrichment of bilirubin-regulated genes in the LRH-1 gene set compared to these negative control ChIP gene sets (\**p*=0.0335, \*\*\**p*=0.0003, by Fisher’s exact test), as would be expected if LRH-1 is a bilirubin receptor. All ChIP gene sets are available as Supplemental Spreadsheets 2 and 3 and references available in methods, briefly AHR ChIP gene set in HepG2 cells was available on the publisher’s website, the PPARα ChIP-seq gene set in HepG2 cells was obtained by contacting authors, PPARγ and SP1 ChIP genes in HepG2 cells were obtained from the ENCODE project. These data suggest bilirubin selectively regulates LRH-1 ChIP-seq target genes, just as frequently as direct AHR and PPARα target genes, two known receptors for bilirubin.

**Supplemental Figure S12:**
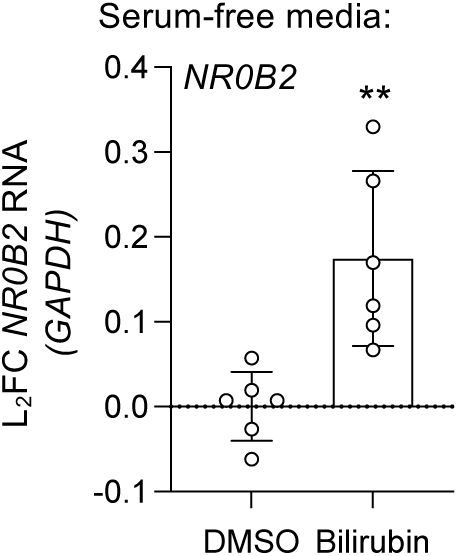
Serum-free media does not alter bilirubin activation of direct LRH-1 target gene NR0B2. RT-qPCR using primers for *NR0B2* from RNA from HepG2 cells expressing endogenous LRH-1 and stably expressing *SLCO1B1* treated with either DMSO vehicle or 50uM bilirubin for 24 hours, error represents SD from the mean, *p=0.0033* by two-tailed unpaired t-test. These data suggest bilirubin can activate NR0B2, a direct target gene of LRH-1, in HepG2 cells in the absence of serum in the media.

**Supplemental Figure S13:**
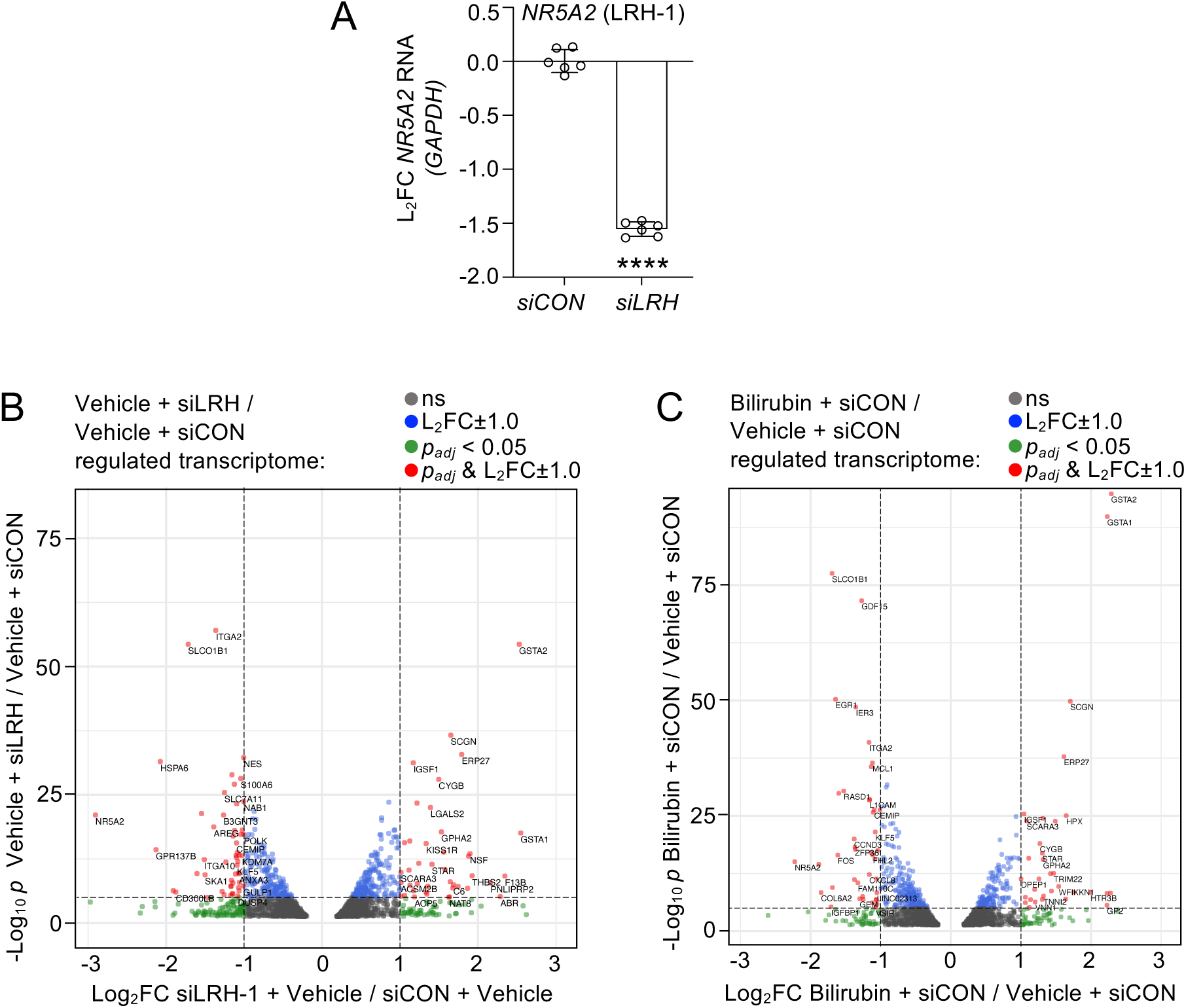
Control transcriptomes suggest siRNA genetic perturbations of LRH-1 regulate gene expression in HepG2 cells, as expected. **A.** RT-qPCR of LRH-1 transcripts (*NR5A2*) regulated by siRNA against LRH-1 (*NR5A2*) after 96 hours transfection of HepG2 cells stably expressing *SLCO1B1*, showing siRNA against LRH-1 decreases significantly decreases LRH-1 transcripts, error represents SD from the mean, \*\*\*\**p*<0.0001 by unpaired, two-way t-test. **B.** Volcano plot of the transcriptome regulated by siRNA against LRH-1 after 96 hours transfection of HepG2 cells stably expressing *SLCO1B1*, showing siRNA against LRH-1 differentially expresses 245 genes (*padj*<0.05 & Log2FC±1), suggesting LRH-1 knockdown by siRNA regulates gene expression in HepG2 cells. **C.** Volcano plot of the 24-hour 50uM bilirubin-regulated transcriptome after 96 hours control siRNA transfection of HepG2 cells stably expressing *SLCO1B1*, suggesting 50uM bilirubin differentially expressed 171 genes (*padj*<0.05 & Log2FC±1) in the presence of control siRNA. These volcano plots contain all the data points from these control transcriptome reactions.

**Supplemental Figure S14.**
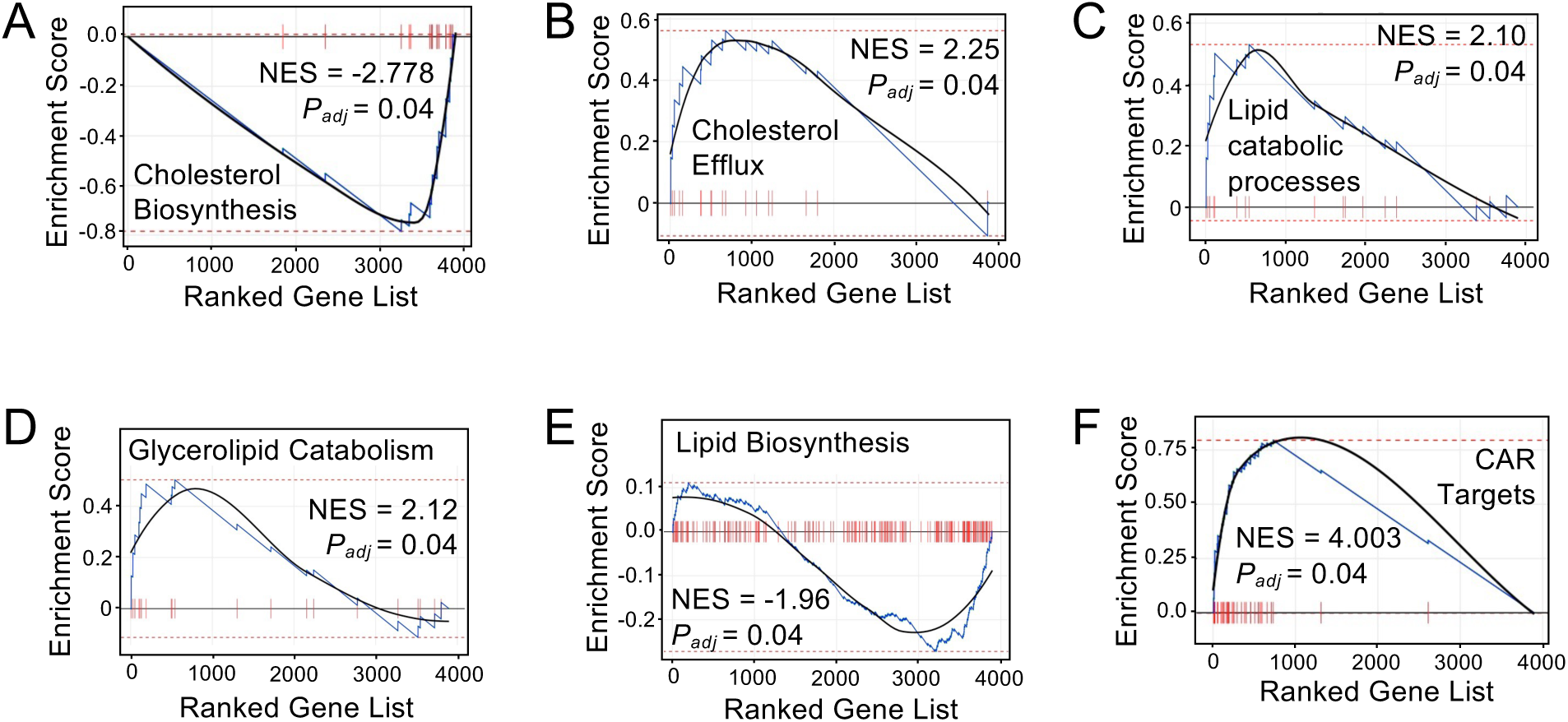
Gene Set Enrichment (GSEA) of the bilirubin-regulated transcriptome in HepG2 cells is enriched in known liver metabolic pathways. Gene Set Enrichment bar code plots of 24 hour, 50uM bilirubin-regulated transcriptomes (bilirubin/vehicle) using the Molecular Signatures database, illustrating **A**. bilirubin-induced de-enrichment of Cholesterol Biosynthesis genes, **B.** bilirubin induced enrichment in Cholesterol Efflux genes, **C**. bilirubin induced enrichment in lipid catabolic processes, **D**. bilirubin induced enrichment in glycerolipid catabolic processes, **E**. bilirubin induced de-enrichment of Lipid Biosynthetic genes, **F**. and bilirubin induced enrichment in nuclear receptor CAR targets. These data suggest bilirubin treatment of HepG2 cells turns off genes associated with cholesterol and lipid synthesis, while turning on genes involved with lipid breakdown and efflux from cells.

**Supplemental Figure S15:**
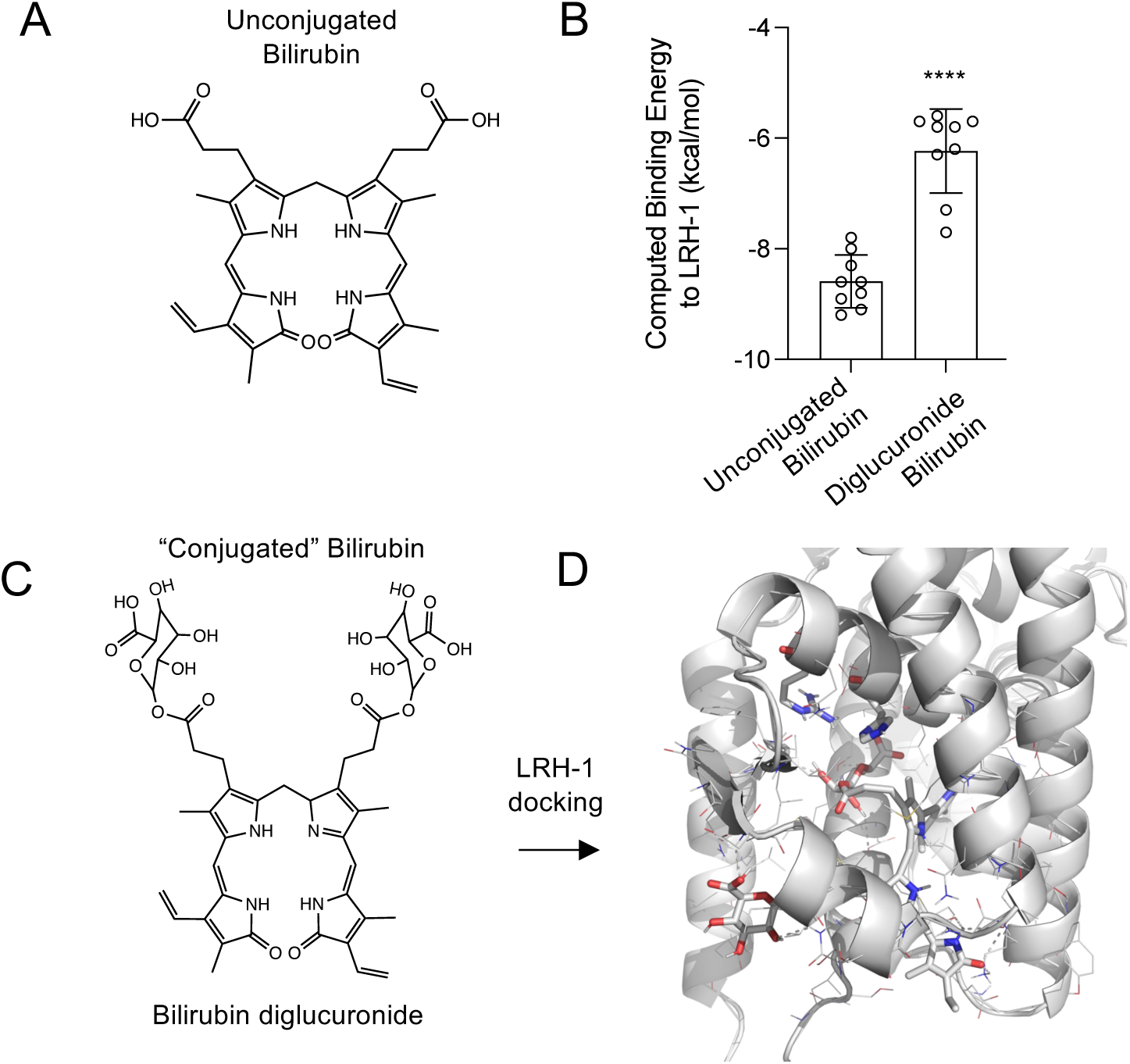
Unconjugated bilirubin is predicted to bind to LRH-1 with lower energy than conjugated bilirubin. **A**. Structure of unconjugated bilirubin used in this study. **B**. Computed binding energies from PyRx rigid body docking to LRH-1 LBD (PDB: 6OQX), showing significantly lower unconjugated bilirubin binding energy compared to conjugated bilirubin *in silico*. **C**. Structure of bilirubin diglucuronide (“conjugated”) and **D.** its PyRx-docked binding position in the canonical LRH-1 ligand binding site. These *in silico* data predict human LRH-1 prefers to bind unconjugated bilirubin, although it remains possible that conjugated bilirubin, or other heme metabolites, may bind LRH-1 in cells. It is important to note that these *in silico* observations cannot easily be tested in the wet lab as unconjugated bilirubin is not commercially available.

## Notes

### Competing Interest Statement

The authors have declared no competing interest.

### Summary of Updates

NIH funding Acknowledgements have been added for National Institutes of Health GM138873.

